# Neuromodulation-induced burst firing in parvalbumin interneurons of the basolateral amygdala mediates transition between fear-associated network and behavioral states

**DOI:** 10.1101/2021.04.19.440525

**Authors:** Xin Fu, Eric Teboul, Jamie Maguire, Jeffrey G. Tasker

## Abstract

Patterned coordination of network activity in the basolateral amygdala (BLA) is important for fear expression. Neuromodulatory systems play an essential role in regulating changes between behavioral states, however the mechanisms underlying the neuromodulatory control of BLA circuits that mediates transitions between brain and behavioral states remain largely unknown. We examined the role of neuromodulation of parvalbumin (PV)-expressing interneurons in the BLA in coordinating network and behavioral states using combined chemogenetics, *ex vivo* patch clamp recordings, and *in vivo* and *ex vivo* local field potential recordings. We show that Gq signaling, whether by the designer receptor, hM3D, α1A adrenoreceptors, or 5-HT2a serotonergic receptors, induces a previously undescribed, highly stereotyped bursting pattern of activity in BLA PV interneurons that generates synchronous bursts of inhibitory postsynaptic currents and phasic firing in the BLA principal neurons. The Gq activation in PV interneurons induced a transition from tonic to phasic firing in the BLA PV neurons and principal neurons and suppressed BLA gamma oscillations in slices and suppressed BLA gamma and potentiated theta power *in vivo*. Gq activation in BLA PV interneurons also facilitated fear memory recall, consistent with previous reports of BLA gamma suppression and theta potentiation during conditioned fear expression. Thus, our data reveal a BLA parvalbumin neuron-specific Gq neuromodulatory mechanism that mediates the transition to a fear-associated network and behavioral state.

## Introduction

Switching between different brain and behavioral states is necessary to adapt to an ever-changing environment. Accompanying brain-state switches are prominent changes in population-level rhythmic and synchronous neural activity (Buzsáki et al., 2012; Lee & Dan, 2012), which can be detected by changes in oscillations of local field potentials and are shaped by the activity of inhibitory interneurons (Bocchio et al., 2017; Buzsáki et al., 2012). Subcortical neuromodulatory systems influencing various cognitive processes, such as arousal, are key mediators of brain state transitions (Lee & Dan, 2012; McCormick et al., 2020), and significantly modulate network rhythmic and synchronous patterning. Increasing evidence indicates that inhibitory interneurons are major targets of neuromodulation and are activated by multiple neuromodulators, such as norepinephrine, serotonin, and acetylcholine, via their cognate G protein-coupled receptors (GPCRs) coupled to Gq signaling pathways (Wester & McBain, 2014; Zagha & McCormick, 2014). Thus, Gq-mediated neuromodulatory regulation of inhibitory interneurons may be critical for orchestrating the changes in neural oscillations that underlie operational state transitions of the brain.

The basolateral amygdala (BLA) acts as a critical node among limbic networks for processing emotionally salient information and patterned BLA network activity has proven central to this process (Courtin et al., 2014; Davis et al., 2017; Kanta et al., 2019; Karalis et al., 2016; Likhtik et al., 2014; Ozawa et al., 2020; Stujenske et al., 2014). Particular importance has been given to BLA theta- and gamma-frequency (~ 2-12 Hz; ~ 40 – 120 Hz, respectively) activity in the dynamic regulation of conditioned fear expression (Courtin et al., 2014; Stujenske et al., 2014). Mechanisms of gamma rhythm generation in other limbic regions point to a coordinating role of parvalbumin-expressing (PV) interneurons (Bartos et al., 2007; Buzsáki et al., 2012; Cardin et al., 2009; Hájos et al., 2004; Sohal et al., 2009), and PV interneurons have been shown to play a critical role in the behavioral expression of fear (Davis et al., 2017; Ozawa et al., 2020). Recent evidence demonstrates a critical role for PV interneurons in generating oscillations in the BLA (Antonoudiou et al., 2021), likely due to their fast-spiking properties and dense perisomatic innervation of hundreds of neighboring principal cells for fast temporal control over synchronous activity (Vereczki et al., 2016; Veres et al., 2017). Further, other reports demonstrate a potential mechanistic role for BLA PV interneurons in coordinating BLA theta oscillations associated with conditioned fear (Davis et al., 2017; Ozawa et al., 2020).

Parvalbumin-expressing interneurons in cortical areas are sensitive to neuromodulatory signals that tune PV cell activity via GPCR activation to execute brain state-dependent behavioral tasks (Garcia-Junco-Clemente et al., 2019; Polack et al., 2013). This provides a potential signaling mechanism for modulatory control of BLA PV cells in brain state transitions and behavior. For example, neuromodulatory signaling in the BLA via norepinephrine plays a well-documented role in promoting conditioned fear states (Giustino & Maren, 2018). Despite this critical role for BLA neuromodulatory signaling in conditioned fear expression, how neuromodulation regulates BLA network rhythms associated with fear states is not known.

Here, we tested the role of Gq neuromodulation of PV cells in controlling network states and regulating conditioned fear expression. Our findings reveal a novel molecular and cellular mechanism for emotional brain state transitions whereby Gq signaling reconfigures BLA PV interneuron activity patterns, which tunes BLA network oscillations and facilitates conditioned fear expression. Stimulation of Gq signaling by the Gq-coupled designer receptor, hM3D, or α1A adrenoreceptor generated phasic firing of BLA PV interneurons and synchronized phasic bursts of IPSCs and phasic firing in BLA principal neurons. Gq activation in BLA PV interneurons suppressed BLA gamma and potentiated theta oscillations in *ex vivo* slices and *in vivo*. Selective rescue of α1A adrenergic signaling or activation of hM3D receptors in BLA PV interneurons in global α1A adrenoreceptor knockout animals enhanced conditioned fear expression. Our findings reveal, therefore, the role played by Gq neuromodulation of BLA PV interneurons in the coordination of BLA population activity that controls network oscillatory states underlying fear memory expression. The generalization of the output to Gq signaling, rather than its dependence on a specific receptor, was unpredicted.

## Results

### Gq activation in BLA PV interneurons stimulates repetitive bursts of IPSCs

To investigate the function of Gq activation in PV interneurons in modulating BLA neural activity, a Cre-dependent AAV virus (AAVdj-DIO-hDLX-hM3D(Gq)-mCherry) was injected bilaterally into the BLA of PV-Cre mice to express Gq-coupled designer receptors exclusively activated by designer drugs (DREADDs) specifically in PV interneurons (Fig. 1A-C). Two weeks after viral delivery of the Gq-DREADDs to BLA PV interneurons, whole-cell voltage clamp recordings were performed in putative BLA principal cells in amygdala slices in the presence of glutamate AMPA-receptor (DNQX, 20 μM) and NMDA-receptor antagonists (APV, 40 μM) to isolate inhibitory postsynaptic currents (IPSCs). Notably, Gq-DREADD activation selectively in PV interneurons with clozapine N-oxide (CNO) (5 μM) induced stereotyped phasic bursts of IPSCs. The repetitive IPSC bursts showed an accelerating intra-burst IPSC frequency, which peaked at higher than 50 Hz and generated a shift in the baseline holding current due to summation (Fig. 1D, E). We observed multiple variants of phasic bursts that had varying amplitudes, durations, and acceleration rates (Fig. 1D), suggesting that the different repetitive bursts were generated by different presynaptic PV interneurons. The PV-mediated repetitive IPSC bursts were of relatively long duration (3.63 ± 0.26 s) and recurred at a low frequency (burst frequency: 0.032 ± 0.001 Hz; inter-burst interval range: 15 to 80 s) (Fig. 1F), and continued for >20 min after CNO was washed from the recording chamber. Consistent with the P/Q-type calcium channel dependence of synaptic output from PV interneurons, and not N-type calcium channel dependence (Chu et al., 2012; Freund & Katona, 2007; Wilson et al., 2001), we found that the CNO-induced IPSC bursts were abolished by the selective P/Q-type calcium channel blocker ω-agatoxin (0.5 μM), but not by the N-type calcium channel blocker ω-conotoxin (1 μM) (Fig. 1H; Supplementary Fig. 1A-D).

**Figure 1.**
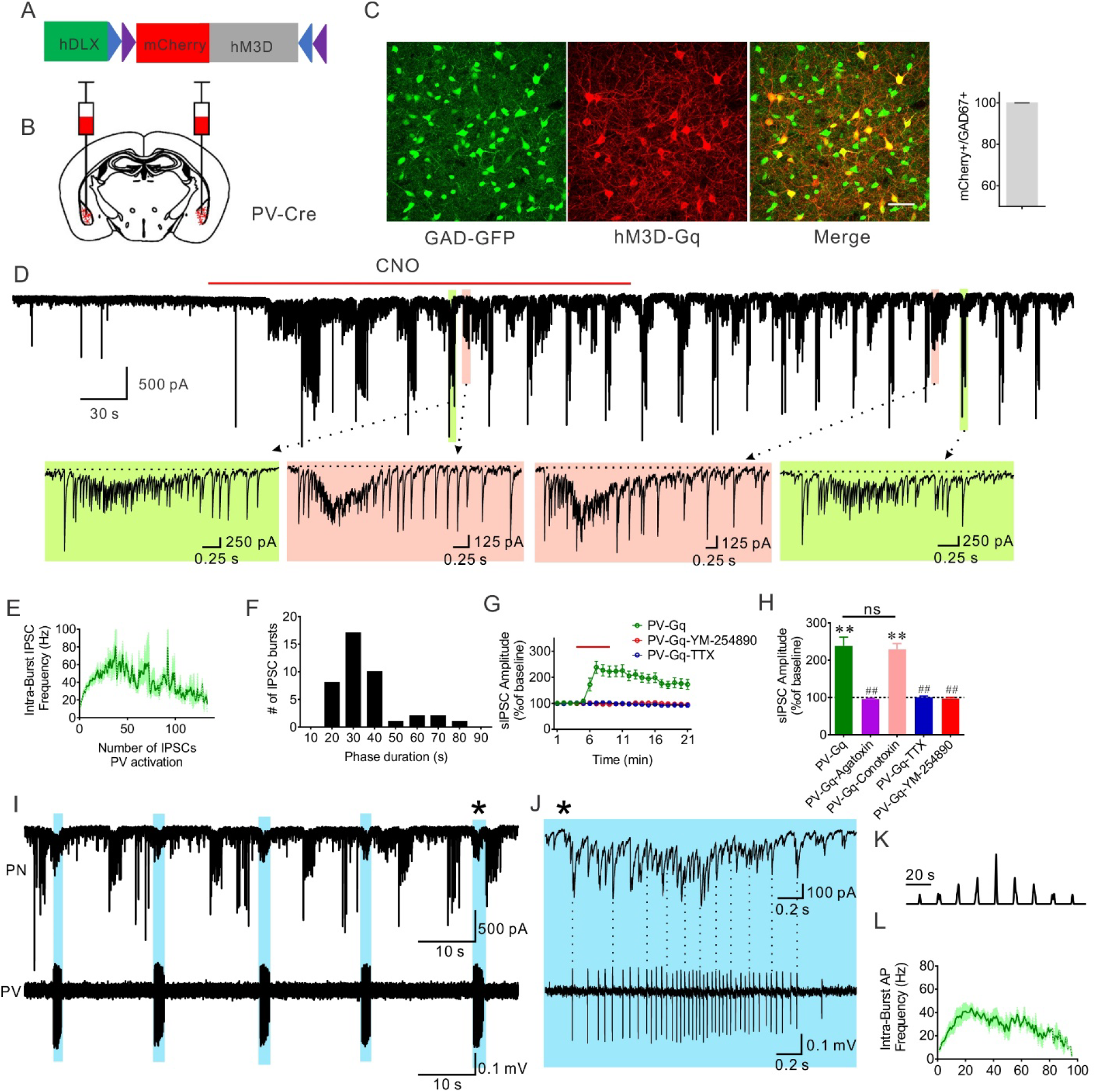
Gq activation in PV interneurons stimulates patterned IPSC bursts in BLA principal neurons. (A, B) Schematic diagrams of bilateral injection of conditional AAV virus expressing a Gq coupled DREADD (hM3D) in the BLA of PV-Cre animals. (C) hM3D(Gq)-mCherry expressed in PV interneurons is co-localized in Gad67-positive GABA neurons (10 sections from 4 animals, 278 cells, ratio=99.81%). (D) A representative recording in a BLA principal neuron showing the generation of phasic IPSC bursts by selective Gq activation in PV interneurons with bath application of CNO. Dashed arrows designate expanded traces of individual bursts to illustrate different stereotyped IPSC bursts (color-coded) at two successive time points. The dashed lines in the expanded traces show the depressed baseline of the holding current due to the summation of high-frequency IPSCs in the bursts. (E) Mean (+/−SEM) of instantaneous intra-burst IPSC frequency over the course of the accelerating IPSC bursts. (F) Distribution of the phase duration of repetitive IPSC bursts; 10-s bins from time −5s to +5s. (41 bursts from 12 cells). (G, H) Time course and mean change in sIPSC amplitude with Gq activation in PV interneurons. The CNO-induced increase in sIPSC amplitude was completely blocked by pre-incubation of the slices with the P/Q-type calcium channel antagonist ω-agatoxin and the sodium channel blocker TTX, and the selective Gα_q/11_ inhibitor YM-254890, but not by the N-type calcium channel antagonist ω-agatoxin (PV-Gq, 12 cells from 5 mice; PV-Gq-Agatoxin, 7 cells from 3 mice; PV-Gq-Conotoxin, 7 cells from 3 mice; PV-Gq-TTX, 8 cells from 4 mice; PV-Gq-YM-254890, 9 cells from 4 mice; Paired *t* test: PV-Gq vs. baseline, p = 0.0001; PV-Gq-Conotoxin vs. baseline, p = 0.0001, **, p < 0.01; One-Way ANOVA, F (4,38) = 22.72, p < 0.0001, Dunnett’s multiple comparisons test, PV-Gq vs. PV-Gq-Agatoxin, p < 0.0001, PV-Gq vs. PV-Gq-Conotoxin, p = 0.98, PV-Gq vs. PV-Gq-TTX, p < 0.0001, PV-Gq vs. PV-Gq-YM-254890, p < 0.0001, ##, p < 0.01, ns, not significant). (I, J) Representative paired recordings showing repetitive IPSC bursts recorded in a BLA principal neuron with whole-cell recording and associated action potential bursts recorded in a PV interneuron with loose-seal recording. Correlated activities are labeled with blue shading and the burst marked with an asterisk was expanded to show the time-locked IPSCs and action potentials. Selected synchronous spikes and IPSCs are designated by vertical dotted lines. (K) Autocorrelation diagram showing the rhythmicity of action potential bursts in the PV neuron shown in I. (L) Mean instantaneous intra-burst frequency (+/− SEM) of action potentials in PV interneurons.

To confirm that the Gq-DREADD-induced IPSC bursts were mediated by the excitation of PV interneurons through Gq signaling, we blocked Gq protein activation with a selective Gα_q/11_ inhibitor YM-254890 (10 μM), which blocks the switch of Gα_q_ from the GDP-to GTP-bound state (Takasaki et al., 2004). We found that the CNO-induced IPSC bursts were completely eliminated with the Gq blocker (Fig. 1G, H). In addition, blocking spiking activity with tetrodotoxin (TTX, 0.5 μM) also abolished the PV Gq-mediated IPSC bursts (Fig. 1G, H), demonstrating the dependence of Gq-induced IPSC bursting in BLA principal neurons on Gq activation in presynaptic PV interneuron somata/dendrites. To determine whether bath application of CNO, which creates a stable drug concentration over minutes, generates a continuous depolarization of PV interneurons that is converted to a phasic synaptic output, we tested whether sustained PV neuron excitation independent of Gq activation is sufficient to generate the repetitive IPSC bursting pattern in principal cells using photoactivation of PV neurons with channelrhodopsin (ChR2). Two weeks after delivery of AAV9-EF1a-DIO-ChR2-mCherry to the BLA of PV-cre mice, continuous photostimulation of PV interneurons with blue light in brain slices failed to generate phasic IPSC bursts in the principal cells, but induced a tonic increase in IPSCs that was phase-locked to the light stimulation (Supplementary Fig. 1E, F). These results together suggest that Gq signaling in PV interneurons is required for the generation of the phasic pattern of inhibitory synaptic output.

The repetitive bursting pattern of IPSCs in BLA principal neurons suggests that presynaptic PV interneurons fire phasic action potential bursts with Gq activation. To directly test this, extracellular loose-seal patch clamp recordings of PV interneurons were performed in slices from PV-Cre mice expressing cre-dependent Gq-DREADDs in the BLA. Similar to the pattern of IPSC bursts in principal neurons, PV cells responded to CNO (5 μM) by firing repetitive accelerating bursts of action potentials that recurred at a low frequency (0.034 ± 0.005 Hz, n = 9 cells) in the presence of DNQX and APV (Fig. 1I-L). Further blockade of fast GABAergic inhibitory synaptic transmission with picrotoxin (50 μM) did not affect the pattern of action potential bursts stimulated by CNO in PV interneurons (n=3, data not shown), suggesting that the bursting activity in PV cells is mediated by an intrinsic rather than circuit mechanism. In paired recordings of synaptically connected PV and principal neurons (n=5 pairs), we found that the action potential bursts in PV neurons were time-locked to one of the subgroups of IPSC bursts in the principal neurons (Fig.1I), confirming that the different variant groups of Gq-induced IPSC bursts in principal neurons were generated by different presynaptic PV interneurons. Moreover, we observed that the action potentials within the bursts of PV cells were time-locked with individual IPSCs in the corresponding IPSC bursts of the principal cells (Fig. 1J). Therefore, Gq activation drives a repetitive bursting pattern of action potentials in BLA PV interneurons to induce patterned bursts of phasic inhibitory synaptic inputs to BLA principal neurons.

Perisomatic PV interneurons innervate hundreds of principal neurons to effectively control spike timing and population-level neural activity (Freund & Katona, 2007; Vereczki et al., 2016; Veres et al., 2017), raising the question whether PV-mediated phasic IPSC bursts are synchronized between BLA principal neurons. To test this, we first examined the synchronization of Gq-induced IPSC bursts with paired recordings from adjacent BLA principal neurons (inter-neuron distance ≤ 40 μm) following CNO activation of hM3D in PV interneurons (Fig. 2A). Interestingly, CNO application induced similar responses in both cells in most of the paired recordings (Fig. 2B). Moreover, we found that the CNO-induced recurrent IPSC bursts were synchronized between each pair at two different levels: 1) the bursts that were synchronized between two cells were synchronous at each repetition, and 2) the individual IPSCs within synchronized bursts were synchronized (Fig. 2B). This is consistent with each specific subtype of IPSC burst being generated by repetitive action potential bursts in a distinct presynaptic PV interneuron. Dividing the total number of PV-mediated IPSC bursts by the number of synchronized bursts, we calculated one BLA principal neuron receives, on average, bursting IPSC inputs from 4.1 presumably different presynaptic PV interneurons, and 67.7% of all the IPSC bursts are synchronized between the pairs of recorded cells (16 recorded pairs from 6 mice) (Fig. 2C).

**Figure 2.**
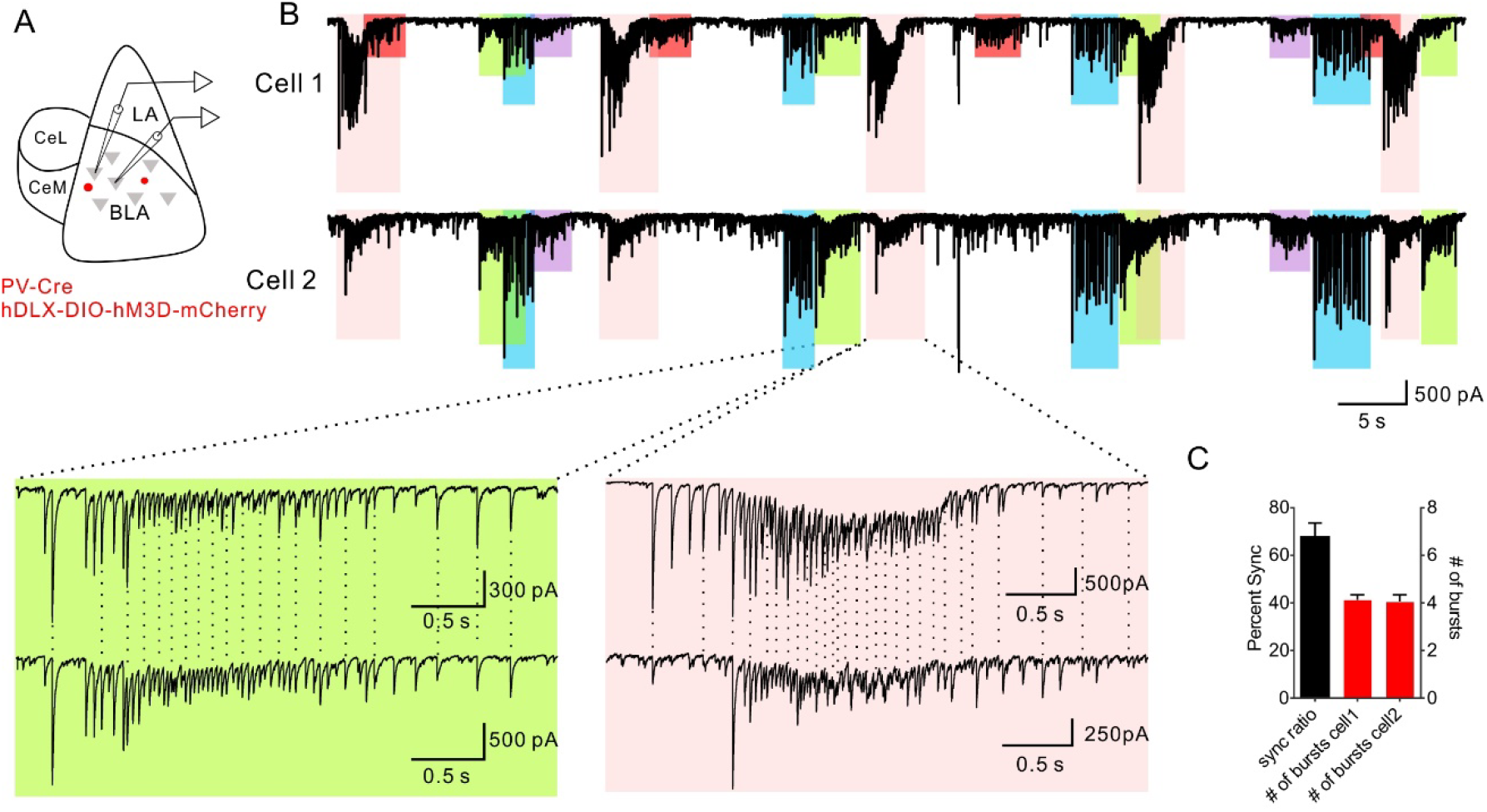
Synchronized Gq-activated IPSC bursts in BLA principal neurons. (A) Schematic showing paradigm of paired recordings of adjacent BLA principal neurons in Gq-DREADD-injected PV-Cre mice. (B) Representative recordings showing synchronized bursts of IPSCs in a pair of BLA principal neurons, Cell 1 and Cell 2. Different colored boxes represent different IPSC bursts repeated over the course of the recording; the same-colored boxes in the two recordings indicate bursts that were synchronized between the two cells. The red-shaded bursts in Cell 1 were not associated with a synchronous IPSC burst in Cell 2. Bottom: Expanded traces of two different, color-coded IPSC bursts from each cell. The dashed vertical lines show the high synchronicity between the two cells of the individual IPSCs that make up each of the bursts. (C) The mean ratio of synchronized bursts to total bursts (Percent Sync), and the total number of different subtypes of bursts (# of bursts) induced in pairs of principal cells by Gq activation of PV interneurons (16 pairs from 6 mice).

### Norepinephrine stimulates PV neuron-mediated repetitive IPSC bursts

Having established the role for Gq signaling in driving repetitive phasic synaptic outputs from PV interneurons in the BLA, we next tested whether native Gq-coupled GPCRs in PV interneurons generate similar bursts of IPSCs in BLA principal neurons. Norepinephrine (NE) is an arousal neuromodulator that is released in the amygdala during stress (McIntyre et al., 2002), and Gq-coupled α1 adrenoreceptors are highly expressed in the BLA (Day et al., 1997). We performed voltage clamp recordings of IPSCs in BLA principal neurons in the presence of glutamate AMPA and NMDA receptor antagonists. NE application (20 – 100 μM) induced a robust, concentration-dependent increase in spontaneous IPSCs that was characterized by a repetitive bursting pattern, which was similar to that elicited by Gq-DREADD activation in PV neurons (Fig. 3A). Following an initial increase in IPSC frequency that inactivated within 35 s to 135 s (mean = 71 s) and was not seen with Gq-DREADD activation in PV neurons, a rhythmic bursting pattern of IPSCs emerged at higher NE concentrations (Fig. 3A-D; Supplementary Fig. 2A, B). The repetitive IPSC bursts occurred at a low frequency (0.028 ± 0.002 Hz), displayed a fast acceleration of intra-burst IPSC frequency that reached peak frequencies > 50 Hz (peak frequency = 50 to 133 Hz), lasted for several seconds (duration, 1.58 to 10.55s, mean = 4.88 ± 0.37s), and gradually tapered off (Fig. 3B-E); these NE-induced IPSC bursts were very similar to the repetitive IPSC bursts stimulated in principal neurons by Gq activation in PV interneurons (see Fig. 1). In some recordings, we observed multiple variants of NE-induced phasic IPSC bursts with differing burst characteristics (Supplementary Fig. 3A), which we interpreted to be generated by multiple presynaptic PV neurons, like the Gq-DREADD-induced IPSC bursts. Individual IPSCs within the NE-induced bursts showed a fast rise time (10-90%: 1.11 ms ± 0.07 ms) and decay time (tau: 18.94 ms ± 1.04 ms), suggesting that they originated from perisomatic inhibitory interneurons, such as cholecystokinin (CCK) or PV basket cells (McGarry & Carter, 2016; Wilson et al., 2001). CCK and PV basket cells can be distinguished by differential expression of voltage-gated calcium channels and CB1 receptors at their synapses (Freund & Katona, 2007; Owen et al., 2013). Double dissociation of NE-induced IPSCs with calcium channel blockers and a cannabinoid receptor agonist showed that while the initial increase in IPSC amplitude was blocked by the N-type calcium channel blocker ω-conotoxin (1 μM) and a CB1 receptor agonist, WIN 55,212-2 (1 μM), the repetitive IPSC bursts were not affected by either ω-conotoxin or WIN 55,212-2, but were selectively blocked by the P/Q-type calcium channel blocker ω-agatoxin (0.5 μM) (Fig. 3A, F, G). Since GABA release from PV interneurons is P/Q-type channel dependent, while GABA release from CCK interneurons is mediated by N-type channels (Freund & Katona, 2007; Wilson et al., 2001) and suppressed by CB1 receptor activation (Neu et al., 2007; Vogel et al., 2016), this indicated that the NE-induced phasic IPSC bursts were generated by GABA release from PV interneuron synapses onto the principal cells. While it abolished the NE-induced IPSC bursts, blocking PV interneuron inputs to principal cells with the P/Q channel antagonist only suppressed the overall increase in IPSC frequency by about 50%, and blocking CCK basket cell inputs with the CB1 receptor agonist failed to reduce the frequency response further (Supplementary Fig. 3B), which suggested that NE also activates other interneuron inputs to the principal cells, possibly from somatostatin cells. The further decrease in the NE-induced increase in IPSC frequency by TTX (Supplementary Fig. 3H-J) suggests that this residual IPSC response to NE is mediated by a combination of NE actions at the somata (TTX-sensitive) and axon terminals (TTX-insensitive) of these unidentified presynaptic interneurons.

**Figure 3.**
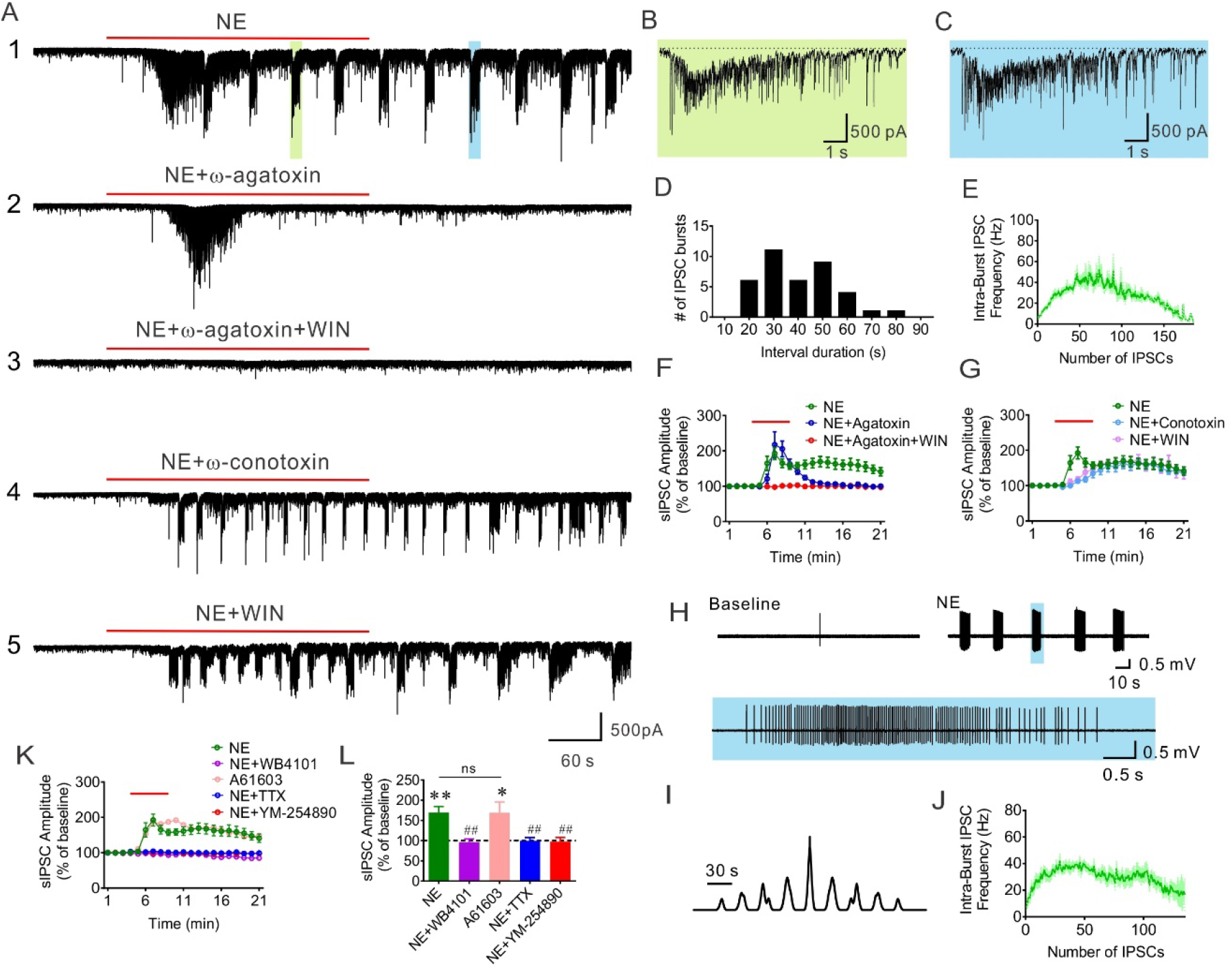
α1A adrenergic receptor activation of PV interneurons generates phasic IPSC bursts in BLA principal cells. (A_1-5_) Representative recordings showing the effect of different treatments on NE-induced repetitive IPSC bursts in the BLA principal neurons. 1. NE application induced phasic IPSC bursts following an initial increase in IPSCs. 2. Inhibiting PV neuron-mediated transmission with the P/Q calcium channel blocker ω-agatoxin selectively blocked the NE-induced repetitive IPSC bursts, but not the initial increase in IPSCs. 3. Co-application of ω-agatoxin and the CB1 receptor agonist WIN 55,212-2 blocked both the initial increase in IPSCs and the repetitive IPSC bursts. 4. Pretreatment of a slice with the N-type calcium channel blocker ω-conotoxin selectively inhibited the initial increase in IPSCs induced by NE, but not the repetitive IPSC bursts. 5. Application of the CB1 receptor agonist WIN 55,212-2 blocked the NE-induced initial IPSC increase, but did not affect the repetitive IPSC bursts. (B, C) Expanded traces of individual IPSC bursts in A1 (indicated with green and blue shading), showing the fast acceleration in the intra-burst IPSC frequency and resulting depression in the baseline holding current (indicated with dashed lines) induced by NE. (D) histogram showing the distribution of phase durations of NE-induced repetitive IPSC bursts (n=38 bursts from 16 cells). (E) Mean instantaneous intra-burst IPSC frequency (+/− SEM) over the course of the NE-stimulated accelerating IPSC bursts. (F) Time course of the effect of blocking P/Q-type calcium channels and activating CB1 receptors on sIPSC amplitude. The NE-induced plateau increase in sIPSC amplitude was blocked by ω-agatoxin (P/Q blocker), while the peak increase, which corresponds to the NE-induced initial increase in IPSCs shown in A2, was unaffected. The ω-agatoxin-insensitive IPSCs were blocked by CB1 receptor activation with WIN 55,212-2, corresponding to the recording in A3. (G) Time course of the effect of blocking N-type calcium channels and activating CB1 receptors on sIPSC amplitude. Both treatments selectively eliminate the NE-induced initial increase in IPSCs, with little effect on NE-induced repetitive IPSC bursts, which corresponds to the recordings in A4 and A5. (H) Representative loose-seal extracellular recording showing the NE-stimulated repetitive bursts of action potentials in a PV interneuron. A burst of action potentials indicated by the blue box was expanded below to show the accelerating intra-burst IPSC frequency. (I) Autocorrelation showing the rhythmicity of NE-induced AP bursts in the PV interneuron shown in H. (J) Mean instantaneous intra-burst action potential frequency (+/− SEM) over the course of PV action potential bursts induced by NE. (K, L) Mean change in sIPSC amplitude over time in response to NE (n=16 cells from 5 mice), NE + α1A receptor antagonist WB4101 (n=7 cells from 4 mice), NE + α1A receptor agonist A61603 (n=10 cells from 4 mice), NE + TTX (n=10 cells from 5 mice), NE + Gq antagonist YM-254890 (n=8 cells from 3 mice); paired *t* tests: NE vs. baseline, p = 0.0003, A61603 vs. baseline, p = 0.023, *, p < 0.05, **, p < 0.01; One-Way ANOVA, F (4, 46) = 6.22, p = 0.0034, Dunnett’s multiple comparisons test, NE vs. NE + WB4101, p = 0.0098, NE vs. A61603, p > 0.99, NE vs. NE + TTX, p = 0.0052, NE vs. NE + YM-254890, P = 0.0078, ##, p < 0.01, ns, not significant; values at time = 13 min were used to compare ω-agatoxin-sensitive IPSCs.

To directly test whether NE activates repetitive bursts of action potentials in PV interneurons, extracellular loose-seal recordings of PV interneurons were performed in slices from PV-Cre animals crossed with Ai14 reporter mice, in which PV interneurons express tdTomato. Application of NE (100 μM) induced repetitive bursts of accelerating action potentials in these neurons that recurred at intervals of tens of seconds (0.03±0.006 Hz, n=7 cells), similar to the NE-induced IPSC bursts in principal neurons (Fig. 3H-J). Note that NE failed to activate PV neurons when they were recorded in the whole-cell current clamp configuration, suggesting that the intracellular signaling mechanism necessary to generate the NE response was washed out with the dialysis of the cytosol. Overall, these data suggest that activation of PV interneurons by NE generates repetitive IPSC bursts that resemble those stimulated by chemogenetic activation of PV interneurons.

To determine whether the NE-induced IPSC bursts are mediated by activation of Gq-coupled adrenoreceptors, we first tested for the adrenoreceptor subtype dependence of the NE-induced increase in IPSCs. Whereas the β adrenoreceptor antagonist propranolol (10 μM) had no effect on the bursts, the NE-induced IPSC bursts were abolished by the broad-spectrum α1 adrenoreceptor antagonist prazosin (10 μM) (Supplementary Fig. 3D-G). The NE-induced increase in IPSCs was also abolished by the α1A adrenoreceptor-selective antagonist WB4101 (1 μM) and mimicked by the α1A adrenoreceptor-selective agonist A61603 (2 μM) (Fig. 3K, L; Supplementary Fig. 3H-J). These results suggest an α1A adrenoreceptor dependence of the NE-induced inhibitory synaptic inputs to BLA principal cells. Consistent with α1A adrenoreceptor signaling through Gq, blocking Gq activation with the Gα_q/11_ inhibitor YM-254890 eliminated all NE-induced IPSCs (Fig. 3K, L; Supplementary Fig. 3H-J). In addition, blocking spiking activity with TTX also inhibited the NE-induced increase in IPSCs (Fig. 3K, L; Supplementary Fig. 3H-J). Therefore, like the chemogenetically-induced phasic IPSC bursts, the NE-induced repetitive IPSC bursts were mediated by Gq activation in presynaptic PV interneurons.

It has been reported that the commercial antibodies against α1 adrenoreceptors are not specific (Jensen et al., 2008). Therefore, to test whether PV interneurons in the BLA express α1A adrenoreceptors, we took advantage of a global α1A adrenoreceptor knockout mouse line (adra1A KO) in which a lacZ gene cassette is placed in frame with the first exon of the adra1A gene, which allows the visualization of α1A adrenoreceptor expression by histochemical staining for β-galactosidase activity with X-gal (Rokosh & Simpson, 2002). We found the adra1A gene to be expressed at high levels in the cortex, hippocampus, amygdala, and hypothalamus (Fig. 4A). In the BLA, we observed a sparse distribution of X-gal-stained cells, which was suggestive of labeled interneurons. To test for the GABA neuron identity of the X-gal-stained cells, we crossed the adra1A KO mouse with a Gad67-eGFP mouse, in which all inhibitory interneurons in the BLA express GFP (Tamamaki et al., 2003). adra1A KO × Gad67-eGFP brains were imaged and analyzed for X-gal and GFP co-staining (Levitsky et al., 2013). To combine the β-galactosidase histochemistry with confocal fluorescence imaging without occlusion of the GFP signal, we performed time-lapse confocal imaging at the same location to track GFP-positive cells before and after the X-gal staining. The sparsely distributed X-gal-labeled cells showed a 98.37% overlap with GFP-labeled GABAergic cells (Fig. 4B, C). We next tested for the expression of α1A adrenoreceptors in PV interneurons by injecting a Cre-dependent AAV virus expressing mCherry into the BLA of PV-Cre mice crossed with the adra1A KO mouse (PV-Cre::adra1A KO) and performed time-lapse imaging of X-gal co-labeling of GFP-positive PV interneurons. Two weeks after virus injection, we observed most of the labeled PV interneurons to be positive for X-gal (84.7 %) (Fig. 4D, E). Together, these data demonstrate that α1A adrenoceptors are selectively expressed in GABAergic interneurons in the BLA, including in most of the PV interneurons.

**Figure 4.**
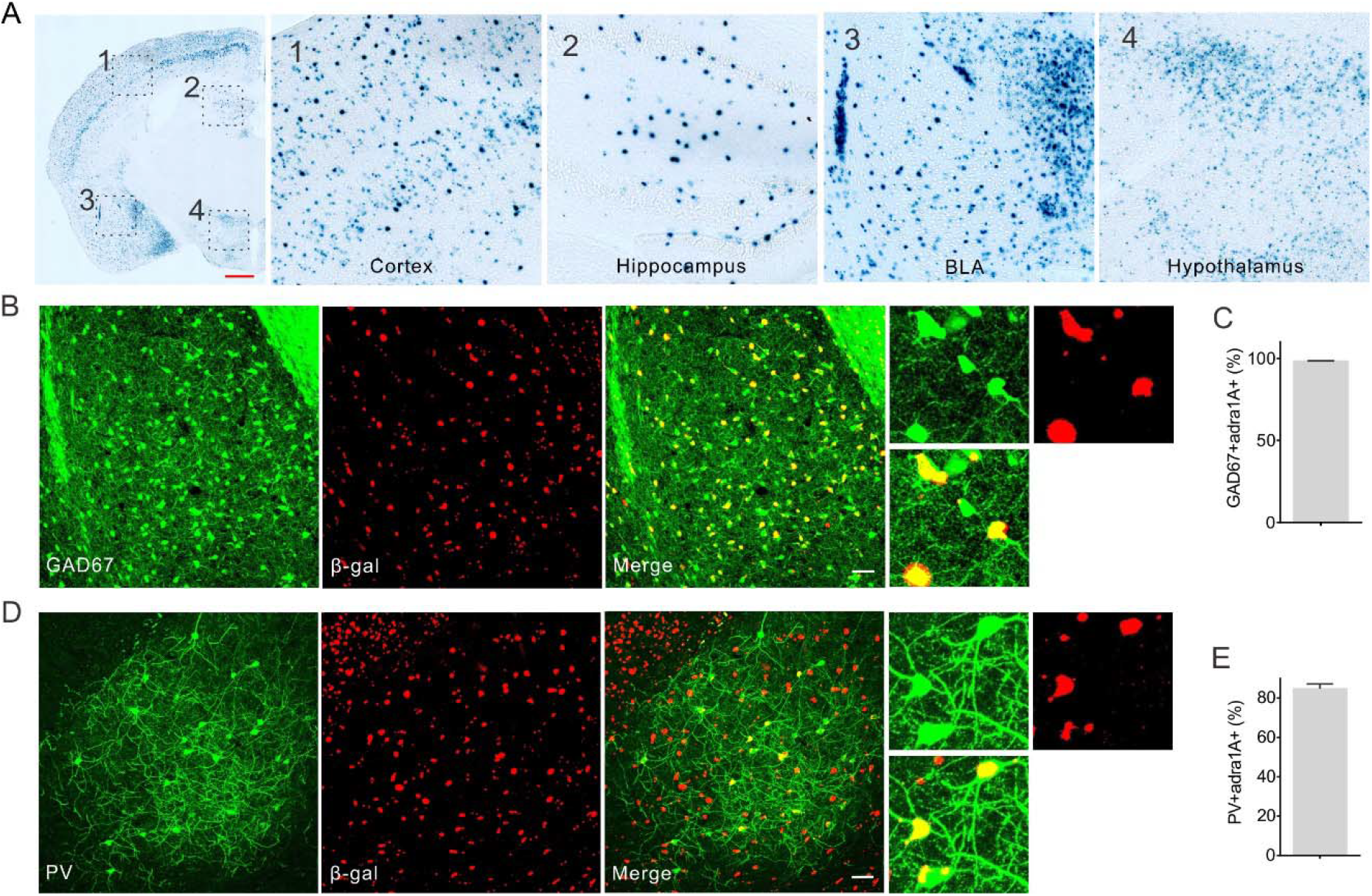
Expression of α1A adrenoreceptors in BLA PV interneurons. (A) Coronal section of the brain showing β-galactosidase staining of α1A adrenoreceptor-expressing cells in the adra1A KO mouse. (1-4) High-magnification images of areas indicated by dashed boxes showing the cortex (1), the hippocampus (2), the BLA (3), and the hypothalamus (4). Scale bar, 500 μm. (B) Colocalization of the β-gal signal with GFP expression in GABA interneurons in the BLA. Scale bar, 50 μm. (C) Percentage of total GAD67-positive cells in the BLA that were positive for β-gal (98.4 ± 0.18%, N=302 cells, 3 sections from 2 animals). (D) Colocalization of the β-gal signal with GFP expression in PV interneurons labelled by injection of Cre-dependent virus in PV-Cre mice. Scale bar, 50 μm. (E) Percentage of GFP-labelled PV interneurons that were positive for β-gal (84.7 ± 2.46%, N= 134 cells, 6 sections from 2 animals).

Multiple neurotransmitter receptors couple to Gα_q/11_ to enhance neuronal excitability, suggesting other neuromodulators may also induce similar phasic IPSC bursts when acting on PV interneurons. Hence, we also tested the effect of serotonin, another neuromodulator that regulates BLA neural circuits (Jiang et al., 2009), on IPSCs in BLA principal neurons. Following the blockade of glutamatergic transmission with DNQX (20 μM) and APV (40 μM) and CCK basket cell-mediated transmission with the CB1 agonist WIN55,212-2 (1 μM), we found that serotonin (100 μM) also induced repetitive bursts of IPSCs that were inhibited by the P/Q calcium channel blocker, ω-agatoxin (0.5 μM), and by an antagonist of the Gq-coupled 5HT2A receptor, MDL 100907 (1 μM) (Supplementary Fig. 4). These data suggest that serotonin also stimulates a phasic synaptic output from PV interneurons, and indicate that Gq activation of PV interneurons serves as a general cellular mechanism for different neuromodulators to regulate BLA neural circuit activity under different emotional states.

### Gq signaling in PV cells alters BLA network states

PV interneurons are critically involved in the generation of gamma-frequency oscillations, which are phase-locked to PV neuron action potentials during tonic high-frequency spiking activity (Bartos et al., 2007; Buzsáki et al., 2012; Hájos et al., 2004). As Gq activation in PV interneurons generated phasic action potentials in PV cells and synchronized repetitive IPSC bursts in principal cells of the BLA, we postulated that Gq-activation of PV cells during tonic activity similarly reconfigures pattern generation among BLA networks and may disrupt potential gamma-frequency network coordination. To address this, we first recorded the spiking responses in BLA PV interneurons and principal cells to PV neuron Gq activation during tonic high-frequency spiking activity.

Loose-seal patch clamp recordings were performed to record spiking activity in hM3D-Gq-transduced PV interneurons using an extracellular potassium concentration that was increased from 2.5 mM to 7.5 mM to increase spontaneous spiking. Strikingly, in all the PV interneurons that displayed spontaneous tonic firing under these conditions (n=7 cells, range: 9.7 to 32.2 Hz, mean: 18.56 ± 3.4 Hz), bath application of CNO (5 μM) transformed the firing pattern of the PV cells from a tonic to a phasic spiking pattern, rather than superimposing the bursts on the baseline tonic spiking activity (Fig. 5A, B). This indicated that Gq activation in PV interneurons causes a switch in the operational mode of BLA PV cells from tonic to phasic activity.

**Figure 5.**
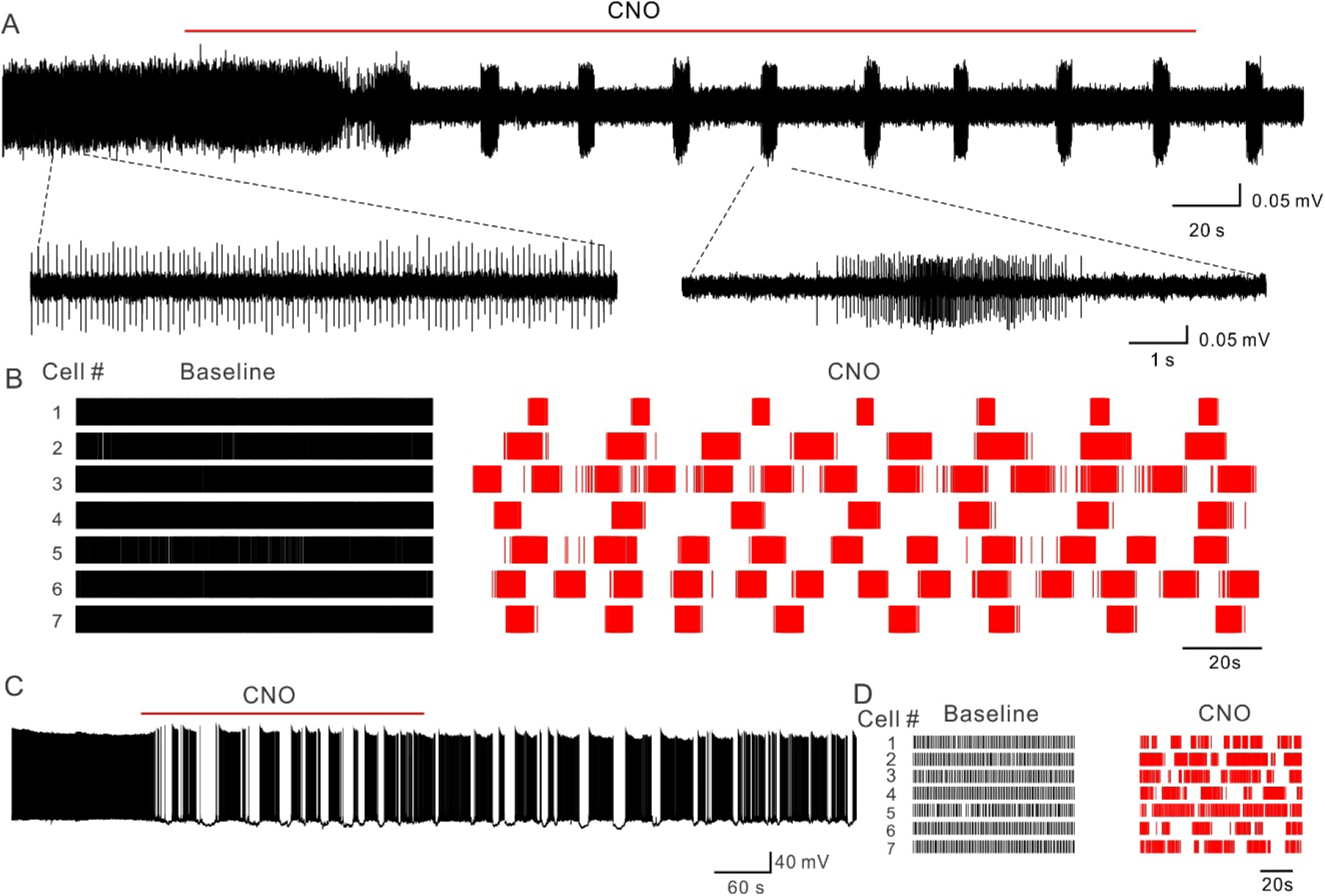
Burst firing in PV interneurons transforms BLA neural activity. (A), A representative recording showing that chemogenetic activation of PV interneurons switches the firing pattern of PV cells from tonic to phasic. Below: Traces were expanded to show the tonic firing in the baseline and burst firing after CNO application. (B), Raster plots of spiking activity in recordings from 7 PV interneurons that show the transformation of tonic spiking to phasic spiking by CNO. (C), A representative current clamp recording from a principal neuron showing that Gq-activation in PV cells transforms the firing pattern of a BLA principal neuron from tonic to phasic. (D) Raster plots of spiking activity in recordings from 7 principal neurons that show the transformation of tonic spiking to phasic spiking by CNO.

To determine the effect of the PV-mediated phasic IPSC bursts on the firing pattern of BLA principal neurons, we recorded from principal neurons in the whole-cell current-clamp recording configuration with a potassium gluconate solution in the recording pipette and activated Gq-DREADDs in PV interneurons with bath application of CNO (5 μM). The membrane potential of recorded principal cells was held above threshold with positive current injection to elicit tonic 2-5 Hz action potential firing, the range of spontaneous spiking activity observed in these cells *in vivo* (Herry et al., 2008; Woodruff & Sah, 2007). Consistent with the change in baseline holding current caused by the accelerating IPSC bursts, activation of repetitive IPSC bursts with CNO application induced prominent oscillatory hyperpolarizations of the membrane potential, shifting the firing pattern of BLA principal cells from tonic to phasic, characterized by slowly oscillating spike bursts at 0.035 ± 0.006Hz (Fig. 5C, D). Together, these data reveal a novel regulatory role for fast spiking interneurons in controlling BLA principal neuron activity patterns in response to Gq activation.

Given the robust influence of Gq signaling on the pattern of BLA PV interneuron activity, we postulated that Gq activation in PV neurons may reconfigure BLA population-level neural activity. To test this, we used a recently developed *ex vivo* slice model for recording locally generated, pharmacologically-induced gamma-frequency BLA network oscillations (30 – 80 Hz; peak frequency = ~30-50 Hz) (Antonoudiou et al., 2021). Local field potentials (LFPs) were recorded in slices from hM3D-Gq-transduced PV-Cre mouse slices of BLA in which inputs from the hippocampus were resected (Supplementary Fig. 5A). Bath application of CNO (5 μM) robustly suppressed slow gamma oscillatory power (Supplementary Fig. 5B-D). We next examined the potential for α1A adrenoreceptor activation to similarly modulate gamma oscillations in resected BLA slices from wild-type mice. Consistently, bath application of 100 μM NE suppressed 30-80 Hz oscillatory power. Further, this effect was inhibited by pretreatment with an α1A adrenoreceptor antagonist, WB4101 (1 μM, Supplementary Fig. 5E-H). Therefore, Gq activation of BLA PV interneurons can reconfigure local network states, suggesting Gq neuromodulation as a potential mechanism for mediating transitions between network and behavioral states.

### Gq signaling in BLA PV cells promotes BLA network state switches *in vivo*

Given the robust influence of Gq signaling on the pattern of BLA PV interneuron activity and local network activity remodeling *ex vivo*, we postulated that Gq activation in PV neurons may also reconfigure BLA population-level neural activity *in vivo*. To test this, we recorded LFPs in the BLA *in vivo* in PV-Cre mice with selective viral expression of hM3D-Gq in BLA PV interneurons (Fig. 6A-C). Gq activation in BLA PV interneurons with intraperitoneal (IP) injection of 5 mg/kg (1 mg/ml) CNO significantly decreased “fast gamma” (70-120 Hz) oscillatory power in the BLA, and produced a trend toward the suppression of “low gamma” (40-70 Hz) power (Fig. 6D-I; Supplementary Fig. 6, 7). Gq activation in BLA PV interneurons also potentiated local theta power, specifically in the “high theta” band (6-12 Hz) (Fig. 6D, G; Supplementary Fig. 6, 7).

**Figure 6.**
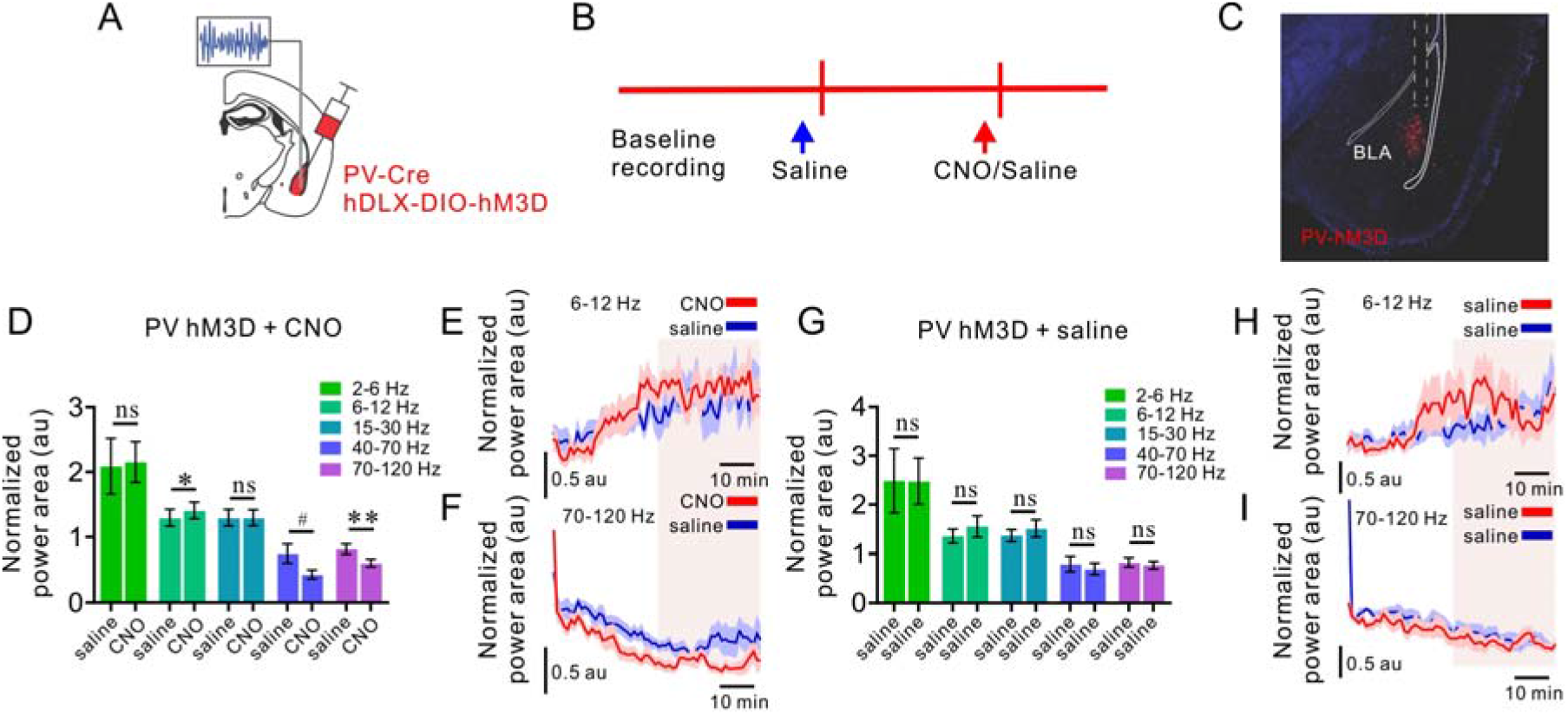
Gq signaling in BLA PV interneurons reconfigures BLA network activity *in vivo*. (A) Schematic illustrating hDLX-DIO-hM3D-mCherry injection and LFP recording strategy in the BLA of PV-Cre mice. (B) Experimental timeline for LFP recordings before (baseline) and after IP injection of saline, followed by saline or CNO. (C) Representative image illustrating hM3D-mCherry expression and LFP recording electrode placement in the BLA (dashed lines). (D) Mean (+/− SEM) normalized power area across treatments (CNO vs saline) and frequency bands. Values from last 30 minutes of recording were used for analysis, illustrated by the highlighted sections in E and F. Two-way ANOVA [treatment × frequency], F (1.318, 7.910) = 5.689, p = 0.0383; Sidaks’s multiple comparisons test: 2-6 Hz, p = 0.9913; 6-12 Hz, p = 0.0440; 15-30 Hz, p > 0.9999; 40-70 Hz, p = 0.0515; 70-120 Hz, p = 0.0095. n = 7. #, p = 0.0515, *, p < 0.05, **, p < 0.01. (E, F) Normalized power area across time for 6-12Hz (E) and 70-120 Hz (F). Values averaged in 1-min bins. Blue/red lines = average power over time, blue/red shaded areas = SEM. (G) Mean (+/− SEM) normalized power area across treatments (Saline 1 vs Saline 2) and frequency bands. Values from last 30 minutes of recording were used for analysis, illustrated by the highlighted sections in H and I. Two-way ANOVA [treatment × frequency], F (1.103, 6.618) = 0.1866, p = 0.7033, n = 7; Two-way ANOVA [treatment], F (1.000, 6.000) = 0.04822, p = 0.8335, n = 7. ns, not significant. (H, I) Normalized power area across time for 6-12 Hz (H) and 70-120 Hz (I). Values averaged in 1-min bins. Colored lines = average power over time, colored shaded areas = SEM. **, p < 0.01, *, p < 0.01, ns = not significant, au = arbitrary units.

**Figure 7.**
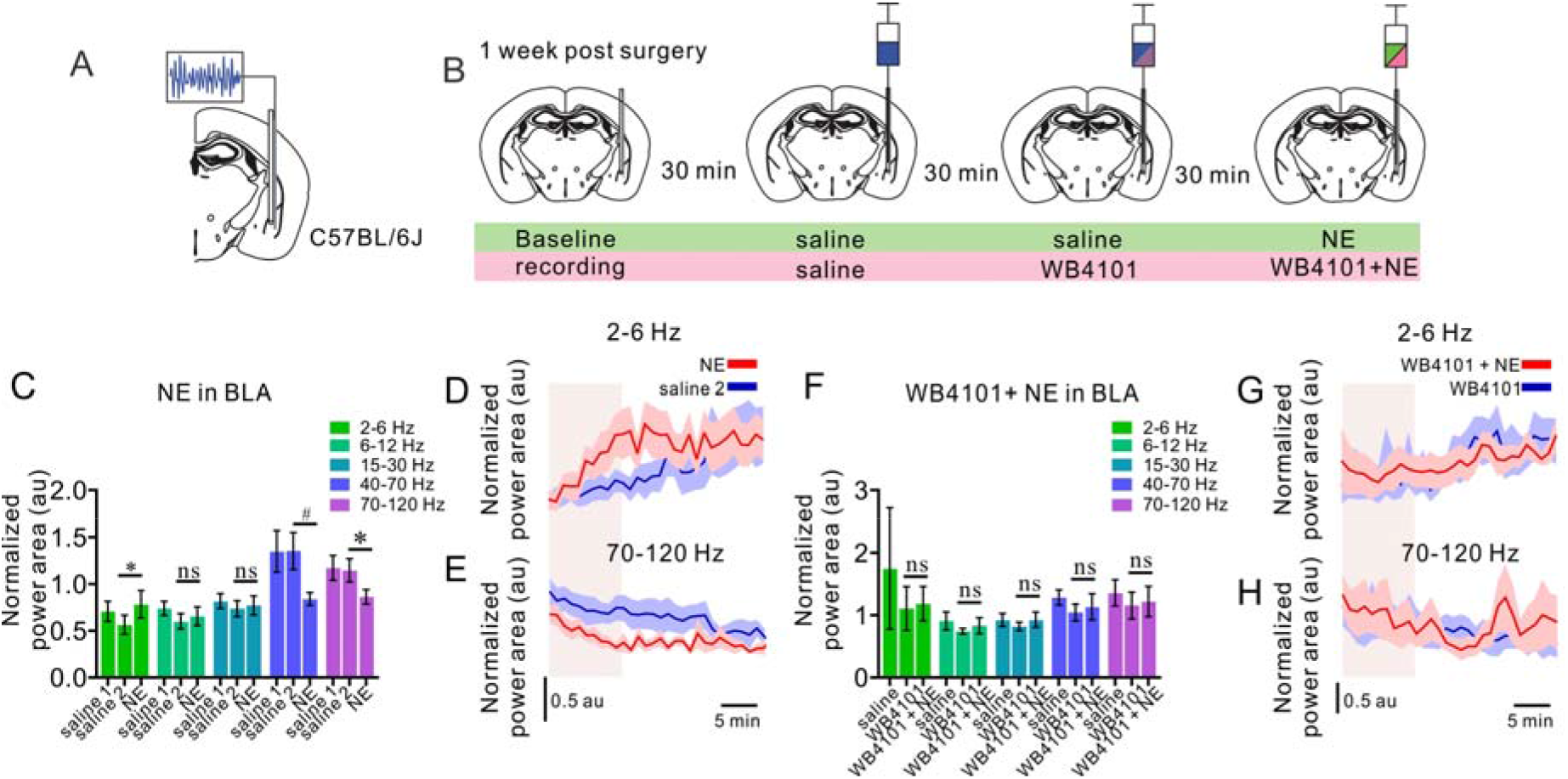
α1A adrenoreceptor signaling dynamically reorganizes BLA network states *in vivo*. (A,B) Schematic illustrating *in vivo* paired intra-BLA drug microinfusion and LFP recording paradigm. (C) Mean (+/− SEM) normalized power area across treatments (Saline1 vs Saline2 vs NE) and frequency bands. Values from first 10 min of each treatment were included for analysis. Two-way ANOVA [treatment × frequency], F (1.657, 11.60) = 5.702, p = 0.0227; Tukey’s multiple comparisons test, 2-6 Hz, p = 0.0310; 6-12 Hz, p = 0.3924; 15-30 Hz, p = 0.5753; 40-70 Hz, p = 0.0794; 70-120 Hz, p = 0.0132. n = 8. #, p = 0.0794, *, p < 0.05. (D,E) Normalized power area across time for 2-6 Hz (D) and 70-120 Hz (E). Values averaged into 1-min bins. Colored lines = average power over time, colored shaded areas = SEM. (F) Mean (+/− SEM) normalized power area across treatments (Saline vs WB4101 vs WB4101+NE) and frequency bands. Values from first 10 min of each treatment were included for analysis. Two-way ANOVA [treatment × frequency], F (1.118, 8.941) = 0.4547, p = 0.5385. Two-way ANOVA [treatment], F (1.818, 14.54) = 0.9321, p = 0.4075, n = 9. ns, not significant. (G,H) Normalized power area across time for 2-6 Hz (G) and 70-120 Hz (H). Values averaged into 1-min bins. Colored lines = average power over time, colored shaded areas = SEM. **, p < 0.01, *, p < 0.01, ns = not significant, au = arbitrary units.

We next tested whether BLA α1A adrenoreceptor activation similarly modulates BLA network oscillations using an *in vivo* neuropharmacological approach (Fig. 7A-B; Supplementary Fig. 8). Intra-BLA infusion of NE (10 mM, 0.3 μL) reduced BLA fast gamma power and potentiated theta power, specifically in the “low theta” (2-6 Hz) band (Fig. 7C-E; Supplementary Fig. 8). These effects were abolished by intra-BLA pretreatment with the α1A-selective adrenoreceptor antagonist WB4101 (10 μM, 0.3 μL) (Fig. 7F-H; Supplementary Fig. 8). Importantly, these data indicate that, like hM3D-Gq activation, α1A adrenoreceptor-dependent noradrenergic signaling in BLA PV interneurons reorganizes network states via an endogenous signaling mechanism *in vivo*.

**Figure 8.**
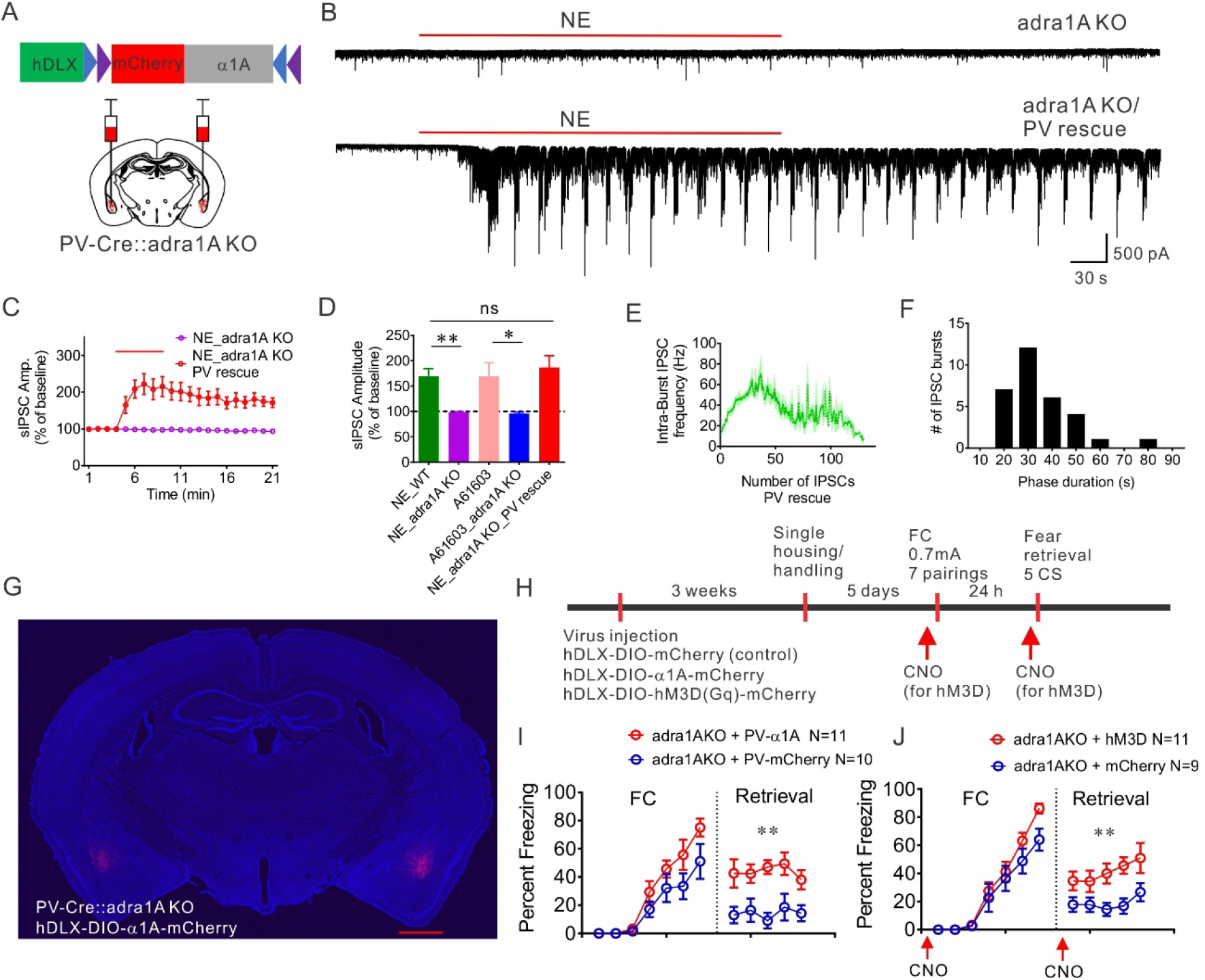
Rescue of α1A adrenoreceptor and Gq signaling in BLA PV interneurons in a global α1A receptor knockout (adra1A KO) mouse facilitates fear memory formation. (A) Schematic diagram of the the bilateral BLA injection sites and the AAV viral vector containing the floxed α1A and mCherry constructs under the control of the GABA neuron-specific hDLX promoter. (B) Representative traces showing the loss of the NE-induced increase in IPSCs in a principal neuron in a BLA slice from an adra1A KO mouse, and the recovery of the NE-induced phasic IPSC bursts in a BLA principal cell following replacement of the α1A adrenoceptors selectively in the PV interneurons. (C) Time course of the NE effect on sIPSC amplitude in adra1A KO mice with or without re-expression of α1A adrenoreceptors in PV interneurons. (D) Mean changes in sIPSC amplitude in BLA principal neurons in response to NE or the α1A-selective agonist A61603 in slices from wild type (WT) and adra1A KO mice. Re-expression of α1A adrenoreceptors in PV interneurons restored the noradrenergic facilitation of sIPSC amplitude in BLA principal neurons. (NE_WT, n=16 cells from 5 mice; NE_adra1A KO, n=9 cells from 4 mice; A61603_WT, n=10 cells from 4 mice; A61603_adra1A KO, n=7 cells from 3 mice; NE_adra1A KO_PV rescue, n = 7 cells from 3 mice) (Unpaired *t* test, A61603_WT vs. A61603_adra1A KO: p = 0.039, *, p < 0.05) (One-Way ANOVA, F (2, 27) = 7.43, p = 0.0025; Dunnett’s multiple comparisons test, NE_WT vs. NE_adra1A KO: p = 0.0046, NE_WT vs. NE_adra1A KO_PV rescue: p = 0.68, **, p < 0.01, ns, not significant). (E) Mean instantaneous intra-burst IPSC frequency over the course of the burst, showing that rescued IPSC bursts display an accelerating intra-burst frequency similar to NE-induced bursts in WT mice. (F) A histogram showing the distribution of the phase duration of NE-induced repetitive IPSC bursts following BLA α1A adrenoreceptor rescue in adra1A KO mice. (G) Representative image showing expression of α1A-mCherry fusion protein after bilateral injections of conditional AAV virus in the BLA of PV-Cre::adra1A KO mice. Scale bar, 1 mm. (H) Experimental timeline for virus injection and fear conditioning, with or without CNO injection. PV-Cre::adra1A KO mice were injected with conditional AAVs expressing Cre-dependent α1A-mCherry or Gq-DREADD to rescue α1A adrenoreceptor mediated Gq signaling in BLA PV interneurons. Control PV-Cre::adra1A KO mice were injected with conditional AAVs expressing only mCherry. For Gq-DREADD rescue experiments, CNO (5 mg/kg) was administrated I.P. 30 min prior to commencement of the fear acquisition and again 30 min prior to commencement of the fear retrieval (red arrows) in both Gq-DREADD and mCherry groups. (I) Replacement of α1A adrenoreceptors selectively in BLA PV interneurons did not have a significant effect on the acquisition (Repeated measures Two-Way ANOVA, F (1, 19) = 3.49, p = 0.078 compared to the control virus-injected PV-Cre::adra1A KO group), but facilitated the retrieval of the fear memory (Repeated measures Two-Way ANOVA, F (1, 19) = 18.47, p = 0.0004). ** p < 0.01 compared to the control virus-injected PV-Cre::adra1A KO group. (J) Rescue of Gq signaling in BLA PV interneurons in adra1A KO mice with excitatory DREADD had no effect on the fear memory acquisition (Repeated measures Two-Way ANOVA, F (1,18) = 2.15, p = 0.16 compared to the control virus injected PV-Cre::adra1A KO group), but enhanced the fear memory retrieval (Repeated measures Two-Way ANOVA, F (1,18) = 11.17, p = 0.0036). ** p < 0.01 compared to control virus injected PV-Cre::adra1A KO group.

### α1A noradrenergic Gq activation of BLA PV interneurons enhances conditioned fear expression

Similar suppression of fast gamma and potentiation of theta oscillatory power among BLA networks have been reported during conditioned fear expression (Stujenske et al., 2014), which suggests a possible BLA PV Gq-mediated neuromodulatory control of fear memory formation. Therefore, we next tested the hypothesis that Gq signaling in BLA PV interneurons promotes conditioned fear expression. To do this, we used a virus-mediated, selective α1A adrenoreceptor re-expression strategy bilaterally in the BLA of a global α1A adrenoreceptor (adra1A) knockout mouse model.

To confirm that selective α1A adrenoreceptor re-expression rescues NE signaling seen with endogenous α1A adrenoreceptors, we first examined NE-induced IPSC bursts using whole-cell recordings in principal neurons in brain slices from global adra1A-knockout mice with and without re-expression of adra1A in BLA PV neurons. To selectively re-express α1A adrenoreceptors in BLA PV interneurons, a Cre-dependent AAV virus expressing α1A and mCherry (AAVdj-hDLX-DIO-α1A-mCherry) was injected bilaterally into the BLA of PV-Cre::adra1A KO mice (Fig. 8A). In slices from adra1A KO mice, NE and the selective α1A adrenoreceptor agonist A61603 failed to generate the phasic IPSC bursts in BLA principal cells seen in slices from wild-type mice (Fig. 8B-D; Supplementary Fig. 9), confirming the specific α1A adrenoreceptor dependence of the NE-induced IPSC burst generation. Re-expression of α1A adrenoreceptors specifically in BLA PV interneurons of adra1A KO mice rescued the NE-induced phasic IPSC bursts in the principal neurons; the NE-induced IPSC bursts showed the characteristic accelerating intra-burst IPSC frequency and low-frequency burst recurrence (0.032 ± 0.002 Hz) seen in principal cells in slices from wild-type mice (Fig. 8A-F).

Having demonstrated the validity of the viral re-expression model, we next investigated the role of α1A noradrenergic activation of BLA PV interneurons in the modulation of conditioned fear expression. Three weeks after α1A adrenoreceptor re-expression in the BLA with bilateral injections of AAVdj-hDlx-DIO-α1A-mCherry into the BLA of PV-Cre::adra1A KO mice (Fig. 8G), the mice were subjected to a standard auditory-cued fear conditioning paradigm (Fig. 8H). Selective rescue of 1A receptors bilaterally in BLA PV interneurons of PV-Cre::adra1A KO mice resulted in enhanced fear memory retrieval on day 2 compared with PV-Cre::adra1A KO mice expressing mCherry alone (Fig. 8I). Note that α1A re-expression in the PV neurons appeared to reverse a loss of fear memory retrieval in PV-Cre::adra1A KO mice. The impairment in fear retrieval was presumably due to Cre and/or control virus expression in the PV neurons and not to the loss of α1A receptors, since α1A receptor deletion in global adra1A KO mice not crossed with the PV-Cre mouse did not show a decrease in fear memory retrieval compared to wild-type mice (data not shown). These data suggest that α1A adrenoreceptor signaling in BLA PV interneurons increases the expression of fear memory.

We next tested the selective activation of hM3D in BLA PV interneurons for changes in fear conditioning to determine the role of PV neuron Gq modulation in fear memory formation. CNO was administered IP 30 min prior to the beginning of the fear acquisition trials on day one, and again 30 min prior to the beginning of the fear memory retrieval trials on day two (Fig. 8H). Similar to the effect of α1A adrenoreceptor rescue on conditioned fear expression, Gq activation in BLA PV interneurons significantly enhanced fear memory retrieval (Fig. 8J). Therefore, our data indicate that Gq signaling in BLA PV interneurons promotes conditioned fear expression.

## Discussion

Our findings reveal a PV cell type-specific neuromodulatory mechanism for BLA network coordination of emotionally salient brain state transitions. Activation of Gq signaling in PV interneurons via hM3D-Gq and α1A adrenoreceptors in PV cells generated a highly stereotyped phasic bursting pattern of activity that drove repetitive, synchronized bursts of IPSCs in postsynaptic BLA principal neurons. Gq-DREADD or α1A activation in BLA PV neurons desynchronized fast gamma and potentiated high theta oscillatory power. Finally, BLA PV Gq signaling enhanced retrieval of an auditory cue-conditioned fear memory, presumably via modulation of BLA network oscillations.

In contrast to the canonical electrophysiological property of PV interneurons to fire sustained high-frequency action potentials upon depolarization, we observed these cells to generate repetitive bursts of action potentials in response to α1A and hM3D receptor-induced Gq activation. The PV neuron phasic bursting was dependent on a postsynaptic intrinsic Gq signaling-dependent mechanism, and not mediated by local circuits or depolarization-induced activation, since phasic activity was not suppressed by blocking ionotropic glutamate and GABA receptors and was not induced with sustained photostimulation of the BLA PV neurons. Therefore, our findings reveal an important role for Gq neuromodulation in changing the operational mode of PV neurons from tonic activation to alternating cycles of activation and inhibition, via an intrinsic mechanism independent of depolarization and fast chemical synaptic transmission. The Gq-dependent cell signaling mechanism responsible for generating the phasic bursting pattern of activation of the BLA PV interneurons is not yet known.

Parvalbumin interneurons are critically involved in the generation of gamma-frequency oscillations in the cortex and hippocampus (Bartos et al., 2007; Buzsáki et al., 2012; Cardin et al., 2009; Hájos et al., 2004; Sohal et al., 2009). Mechanistic studies in these structures indicate that tonic PV neuron activity provides inhibitory signaling to principal neurons that drives the local gamma rhythm, demonstrated by phase locking of PV neuron activity to the gamma cycle (Bartos et al., 2007; Hájos et al., 2004) and by optogenetic activation and inhibition of PV neuron activity (Antonoudiou et al., 2021; Cardin et al., 2009; Sohal et al., 2009). Like in cortical structures, PV interneurons in the BLA densely target the perisomatic region of principal neurons to modulate amygdalar output (Davis et al., 2017; Ozawa et al., 2020; Vereczki et al., 2016; Veres et al., 2017). Therefore, altering BLA PV activity patterns is likely to have a robust effect on BLA network oscillations. We found that Gq activation in BLA PV neurons transformed the tonic activity pattern of the PV neurons into a bursting pattern of action potentials and inhibitory synaptic inputs to BLA principal cells, which induced a phasic pattern of activation of the principal cells. Consistent with tonic PV neuron activity promoting gamma oscillations, transforming PV neuron tonic activity to phasic activity by either chemogenetic manipulation or α1A adrenoreceptor activation decreased gamma oscillatory power locally in the BLA both *ex vivo* and *in vivo*.

Previous work also demonstrated the capacity of BLA PV interneurons to modulate local theta frequency network activity (Davis et al., 2017; Ozawa et al., 2020), and that BLA theta rhythms play a central role in facilitating conditioned fear behavior (Stujenske et al., 2014; Likhtik et al., 2014; Davis et al., 2017; Ozawa et al., 2020). Similarly, our data reveal that both hM3D and α1A receptor activation are sufficient to potentiate theta oscillatory activity, though at separate within-theta frequency bands. We show that while BLA α1A adrenoreceptor activation facilitates a preferential increase at low theta frequencies (2-6 Hz), BLA PV-specific hM3D signaling selectively potentiates the faster high theta frequencies (6-12 Hz). The mechanisms governing the differences between the two manipulations on theta band power are not immediately clear, however differences between α1A adrenoreceptor and hM3D signaling dynamics may play a role. Additionally, selective Gq signaling in BLA PV interneurons may only be activating one aspect of the α1A-induced theta-frequency oscillatory machinery. α1A signaling in other interneuron types in the BLA may work in tandem with local PV interneurons to shape the local field potential to preferentially oscillate at the slower 2-6 Hz. Our finding that NE activates inhibitory synaptic inputs to the BLA principal cells from other GABA interneuron types supports this possibility.

Parvalbumin interneurons in the BLA target the perisomatic region of hundreds of postsynaptic principal neurons to control BLA output. Paired recordings from neighboring principal neurons revealed that a large percentage of the PV-mediated bursts of IPSCs are synchronized. In addition, PV-mediated IPSC bursts transformed the firing pattern of local principal neurons from a tonic to a slow phasic activity. However, other than the disruption of gamma wave generation, it is not known whether the slow synchronous phasic PV neuron inhibitory inputs to BLA principal neurons play a physiological role in pattern generation among BLA circuits. The potentiation of theta power in the BLA LFP suggests that the bursting activity of individual presynaptic PV neurons is staggered to give rise to alternating synchronous IPSC bursts at theta frequency in the principal neurons.

Given the influence of BLA noradrenergic signaling on promoting fear states, we examined what significance the PV neuron α1A adrenoreceptor-mediated reorganization of BLA network activity has in promoting fear states. Genetic re-expression of α1A adrenoreceptors and hM3D-Gq activation in PV interneurons bilaterally in the BLA of adra1A global knockout mice enhanced conditioned fear expression. Therefore, the Gq-mediated transition in BLA PV interneuron patterned output is important for promoting BLA fear state transitions. Conditioned fear expression is associated with BLA potentiation of theta and suppression of fast gamma oscillatory power (Stujenske et al., 2014). Our data demonstrating that Gq signaling via α1A adrenoreceptor and hM3D receptor activation in BLA PV interneurons potentiates local theta and desynchronizes fast gamma network activity reveal a potential critical role for Gq neuromodulation of BLA PV cells in the expression of fear memory.

Thousands of G protein-coupled receptors converge onto four main classes of known Gα proteins (Pierce et al., 2002). In addition to α1A adrenoreceptor-mediated Gq activation of PV interneurons, we also found that Gq-coupled 5-HT2A serotonergic receptors generated a similar pattern of repetitive bursts of IPSCs in BLA principal neurons. Others have reported that acetylcholine also drives repetitive IPSC bursts in pyramidal neurons of the frontal cortex, possibly through the activation of Gq-coupled M1 muscarinic receptors (Kondo & Kawaguchi, 2001), although whether this response is also mediated by PV interneuron activation is not known. Notably, PV interneurons in the hippocampus and BLA stimulated by CCK activation of Gi/o-coupled CCK_B_ receptors (Chung & Moore, 2007; Chung & Moore, 2009; Földy et al., 2007; Lee et al., 2011) generate tonic IPSCs in principal neurons. Thus, different neurotransmitters acting on their cognate Gq-coupled receptors have a similar effect on neural activity through activation of the same signaling pathway, whereas the neuromodulatory excitation of PV interneurons via different GPCR signaling mechanisms induces distinct output patterns. If the patterned bursts of IPSCs is a generalized PV neuron output mode following activation of Gq-coupled receptors by different neuromodulators, then the specificity of the neuromodulatory regulation must lie in other spatio-temporal properties of the neuromodulatory inputs, such as the timing of the inputs and the somato-dendritic distribution of the synapses or synaptic receptors. Then again, the convergence of disparate neuromodulatory inputs onto the Gq signaling pathway to generate a similar patterned PV neuron output may represent a redundancy of neuromodulatory systems to achieve a common salient network outcome, the desynchronization of BLA circuits in the gamma frequency band, under different environmental conditions. Just as different representations of intrinsic membrane ionic currents in individual neurons and synaptic strengths in neural circuits can be tuned homeostatically to achieve target network outputs (e.g., Prinz and Marder, 2004), so may neuromodulatory inputs be activated variably, alone or in combination, to arrive at a particular desired brain state.

In conclusion, our findings provide a cell type-specific neuromodulatory mechanism for BLA network-driven transitions in fear-associated brain states. Thus, during conditioned fear recall, BLA PV neurons transition from a tonic, high-frequency pattern to a phasic bursting pattern of activity in response to Gq activation, which suppresses gamma and potentiates theta oscillatory activity in the BLA. This BLA PV neuron-mediated transition in brain network state facilitates the expression of conditioned fear memory.

## Acknowledgments

We thank Dr. Laura Harrison for her invaluable input to the study. We also thank India Pursell and Hunter Bernstein for their technical assistance. This work was supported by NIH grants R01MH104373 and R01MH119283 (JGT and XF) and R01AA026256, R01NS105628 and R01NS102937 (JLM and ET).

## Author contributions

Conceptualization, J.G.T, X.F., E.T. and J.M.; Investigation, X.F. and E.T.; Writing – Original Draft, X.F. and E.T.; Writing – Review and Editing, J.G.T. and J.M.; Funding acquisition, J.G.T. and J.M.

## Methods

### Animals

Mice were maintained in an AALAC-approved, temperature-controlled animal facility on a 12-h light/dark cycle with food and water provided *ad libitum*. C57BL/6J (Cat. 000664), PV-Cre (Cat. 017320), and adra1A KO mice (Cat. 005039) were purchased from Jackson Laboratories and bred in-house to establish colonies. Heterozygous GAD67-eGFP mice were purchased from Riken BioResource center (Tamamaki et al., 2003) and back-crossed for more than 5 generations with wildtype C57BL/6 mice. All procedures were approved by the Tulane and Tufts University Institutional Animal Care and Use Committees and were conducted in accordance with Public Health Service guidelines for the use of animals in research.

### Stereotaxic Surgery

#### Intracerebral virus injections

##### Intracellular recording

Four- to 6-week-old male mice were anesthetized by intraperitoneal injection of ketamine/xylazine (100 mg/kg) and placed in a stereotaxic frame (Narishige, SR-6N). The scalp was cut along the midline and the skull was exposed. Two burr holes were perforated above the BLA with a Foredom drill (HP4-917). Mice were then injected bilaterally with 350 nl of virus into the BLA (AP: −0.8, ML: 3.05, DV: 4.4) through a 33-gauge Hamilton syringe (10 μl) connected to a micropump (World Precision Instrument, UMP-2) and controller (Micro4) at a flow rate of 100 nl per minute. After waiting for 5 minutes following virus injection to minimize virus spread up the needle track, the injection needle was then slowly retracted from the brain. After surgery, the scalp was sealed with Vetbound, a triple antibiotic ointment was applied, and an analgesic (Buprenorphine, 0.05 mg/kg) was injected IP.

##### Extracellular recording and drug application

Eight- to 10-week-old mice were anesthetized with IP ketamine/Xylazine (100mg/kg; 10mg/kg) and placed in a mouse stereotaxic frame (World Precision Instruments, 502600) over a warm heating pad. Lacri-lube was placed over the subjects’ eyes, and slow-release buprenorphine (Buprenorphine SR-LAB, 0.5 mg/kg) was administered subcutaneously for post-operative analgesia. The scalp was shaved, cleaned with ethanol and betadine (3×), then cut along the midline to expose the skull. The skull was leveled, then manually drilled through above the BLA (relative to bregma: AP −1.35, ML ± 3.3) using a sterile 27G syringe needle. In dual electrode and intra-BLA cannula implantation surgeries, a second hole was manually drilled at AP −3.05, ML ± 3.3. For viral injections, a pulled glass pipette was used to inject 350nL of virus into the BLA (relative to bregma: AP −1.35, ML ± 3.3, DV −5.1) at a flow rate of 100nL per minute using positive pressure from a 10mL syringe. After waiting 5 minutes following injection to minimize viral spread up the needle track, the injection needle was then slowly retracted from the brain. For LFP electrode implants, prefabricated headmounts (Pinnacle Technology Inc, #8201) were mounted to the skull using stainless steel screws that acted as ground and reference, while trying to avoid damaging the cortex. The BLA depth electrode (PFA-coated stainless steel, A-M Systems) was implanted using the same coordinates for virus injection (relative to bregma: AP −1.35, ML ± 3.3, DV −5.1). For paired intra-BLA cannula and recording implantation, the recording equipment was implanted and fixed to the skull as described. A fabricated guide cannula was inserted over the BLA, through the posterior drilled hole (relative to bregma: AP −3.05, ML ± 3.3, relative to nearby skull: −3.08) at a 69-degree angle (Supplementary Fig. 10). The cannula was fixed in place with dental cement (A-M Systems, #525000 and #52600) before releasing from the stereotaxic arm. After surgery, the scalp was either 1) sealed with Vetbond and triple antibiotic ointment was applied, or 2) covered along with the headmount/guide cannula with dental cement and allowed to cure before removing the animal from the stereotaxic frame. The mice were then taken out of the stereotaxic frame and placed in a heated recovery chamber until conscious.

### AAV virus development

For cloning of Cre-dependent hDlx AAV virus, we amplified the hM3D-mCherry and mCherry coding sequence from the plasmid pAAV-hSyn-DIO-hM3D(Gq)-mCherry (a gift from Bryan Roth, Addgene # 44361) (Krashes et al., 2011) and pAAV-hSyn-DIO-mCherry (also a gift from Bryan Roth, Addgene # 50459) and cloned it into a pAAV-hDlx-Flex-GFP vector backbone (a gift from Gordon Fishell, Addgene #83895) (Dimidschstein et al., 2016) at AccI and NheI cloning sites. The coding sequence of adra1A was synthesized from Bio Basic Inc. and cloned into a pAAV-hDlx-Flex backbone. AAV virus from Vigene Biosciences Company was further packaged in AAVdj serotype. All hDlx AAV viruses were diluted to the range of 10^11^ to 10^12^ viral genome per ml with virus dilution buffer containing 350 mM NaCl and 5% D-Sorbitol in PBS.

### Fear conditioning and retrieval

3 weeks after virus injection, mice were single housed and handled for more than 5 days before undergoing a fear conditioning paradigm with the Video Fear Conditioning System in a sound attenuated chamber (MED Associates, Inc.). Each chamber is equipped with a metal stainless-steel grid connected to a shock generator (ENV414S Aversive Stimulator). The fear conditioning paradigm consisted of 7 exposures to a continuous tone (7 kHz, 80 db, 30 s duration) as the conditioned stimulus (CS), each of which was co-terminated with an unconditioned aversive stimulus (US) consisting of electric foot shocks (0.7 mA, 2 s duration). The CS-US stimuli were presented at a randomized intertrial interval (ITI, 30-180 s, average = 110 s) in one context, context A. Twenty-four hours later, on day 2, mice were tested for fear retention in a different context, context B, with a planar floor and a black plastic hinged A-frame insert. During fear memory retrieval, five presentations of CS alone were delivered with an inter-stimulus interval of 60 s. Behavior was recorded with an infrared camera and analyzed with Video Freeze software (Med Associates, Inc.). Mice were considered to be exhibiting freezing behavior if no movement other than respiration was detected for ≥ 2 s. Chambers were cleaned with either 70% ethanol or 3% acetic acid before each session of fear conditioning or fear memory retrieval.

### Histology

#### Perfusion and cryosectioning

two weeks after AAV virus injection, adra1A KO, PV-Cre, and PV-Cre::adra1A KO were deeply anesthetized with ketamine/xylazine (300mg/kg) and perfused transcardially with 10 ml ice-cold PBS (pH 7.4) followed by 20 ml 4% paraformaldehyde (PFA) in PBS. Brains were dissected out, postfixed for 3 hours in 4% PFA in PBS, and cryopreserved with 30% sucrose in PBS for 24 hours at 4°C until the brain sunk to the bottom of the container. Coronal sections (45 μm) were cut on a cryostat (Leica) and harvested in 24-well plates filled with PBS.

#### Confocal imaging

Sections from virus-injected PV-Cre and PV-Cre::adra1A KO mice containing the BLA were selected, rinsed with PBS (3 × 5 mins), and mounted on gel-coated slides. Confocal images were acquired with a Nikon A1 confocal microscope to capture the DAPI (excitation 405 nm, emission 450 nm), GFP (excitation 488 nm, emission 525 nm), and mCherry (excitation 561 nm, emission 595 nm) signals. For the analysis of colocalization, z stack pictures were imaged under a 40x oil-immersion objective with a step increment of 1.5 μm. The number of BLA cells containing colocalized markers were first quantified from merged maximal intensity images from different channels, and then confirmed in z-stack images with ImageJ (NIH).

#### X-gal staining

Sections from adra1A KO mice were rinsed with PBS (3 × 5 min) and incubated in a β-gal staining solution (Roche, Ref # 11828673001) overnight. After β-gal staining, sections were then rinsed in PBS (3 × 5 min), mounted on gel-coated glass slides, coverslipped with Permount mounting medium (Fisher Scientific), and allowed to air dry. Bright-field imaging was performed in a Zeiss Axio Scanner and processed and analyzed with ImageJ (NIH). For fluorescence confocal imaging, brain sections were rinsed with PBS (3 × 5 min), mounted on gel coated coverslips, and then imaged first on the confocal microscope (Nikon A1) before incubating them in the β-gal staining solution. After staining with X-gal, the slices were rinsed with PBS (3 × 5 mins) and the same regions previously imaged using fluorescence confocal imaging were then re-imaged for the X-gal signal with excitation and emission wavelengths of 638 ad 700 nm (Levitsky et al., 2013), respectively. The images were then processed and quantified with ImageJ software to determine the ratio of β-gal-positive cells to fluorescent cells with the same procedure described above.

### Brain slice electrophysiology

#### Patch Clamp

##### Slice preparation

Coronal brain slices containing the BLA were collected from male mice for *ex vivo* electrophysiological recordings. Mice (6 to 9 weeks) were decapitated in a restraining plastic cone (DecapiCone, Braintree Scientific) and the brains were dissected and immersed in ice-cold, oxygenated cutting solution containing the following (in mM): 252 sucrose, 2.5 KCl, 26 NaHCO_3_, 1 CaCl_2_, 5 MgCl_2_, 1.25 NaH_2_PO_4_, 10 glucose. The brains were trimmed and glued to the chuck of a Leica VT-1200 vibratome (Leica Microsystems) and 300 μm-thick coronal slices were sectioned. Slices were transferred to a holding chamber containing oxygenated recording artificial cerebrospinal fluid (aCSF) containing (in mM): 126 NaCl, 2.5 KCl, 1.25 NaH_2_PO_4_, 1.3 MgCl_2_, 2.5 CaCl_2_, 26 NaHCO_3_, and 10 glucose. They were maintained in the holding chamber at 34°C for 30 min before decreasing the chamber temperature to ~20°C.

##### Patch clamp recording

Slices were bisected down the midline and hemi-slices were transferred one-at-a-time from the holding chamber to a submerged recording chamber mounted on the fixed stage of an Olympus BX51WI fluorescence microscope equipped with differential interference contrast (DIC) illumination. The slices in the recording chamber were continuously perfused at a rate of 2 ml/min with recording aCSF maintained at 32-34°C and continuously aerated with 95% O_2_/5% CO_2_. Whole-cell patch clamp recordings were performed in putative principal neurons in the BLA. Glass pipettes with a resistance of 1.6-2.5 MΩ were pulled from borosilicate glass (ID 1.2mm, OD 1.65mm) on a horizontal puller (Sutter P-97) and filled with an intracellular patch solution containing (in mM): 110 CsCl, 30 potassium gluconate, 1.1 EGTA, 10 HEPES, 0.1 CaCl_2_, 4 Mg-ATP, 0.3 Na-GTP, 4 QX-314; pH was adjusted to 7.25 with CsOH and the solution had a final osmolarity of ~ 290 mOsm. DNQX, APV, TTX, Prazosin, Propranolol, WB 4101, A61603, CNO, and NE were delivered at the concentrations indicated via the perfusion bath. Slices were pre-incubated in aCSF containing ω-agatoxin (0.5 μM, 30 min), ω-conotoxin (0.5 μM, 30 min), or YM 254890 (10 μM, 20 min) to block P/Q-type calcium channels, N-type calcium channels and Gα_q/11_ activity, respectively (Takasaki et al., 2004; Owen et al., 2013). The same solution as that used for the aCSF was used in the patch pipettes 1.6-2.5 MΩ) for loose-seal extracellular recording of action potentials, which were performed in the I = 0 mode on the patch clamp amplifier. For current clamp recordings, an intracellular patch solution was used containing (in mM): 130 potassium gluconate, 10 HEPES, 10 phosphocreatine Na_2_, 4 Mg-ATP, 0.4 Na-GTP, 5 KCl, 0.6 EGTA; pH was adjusted to 7.25 with KOH and the solution had a final osmolarity of ~ 290 mOsm. Series resistance was normally below 10 MΩ immediately after break through the membrane and was continuously monitored. Cells were discarded when the series resistance exceeded 20 MΩ. Data were acquired using a Multiclamp 700B amplifier, a Digidata 1440A analog/digital interface, and pClamp 10 software (Molecular Devices). Recordings were filtered at 2 KHz for IPSC recordings and at 10 KHz for action potential recordings and sampled at 50 KHZ. Data were analyzed with MiniAnalysis software (SynaptoSoft, NJ) and Clampfit 10 (Molecular Devices). Statistical comparisons were conducted with a paired or unpaired Students’s *t* test or with a one- or two-way ANOVA followed by a *post hoc* Tukey’s test as appropriate (p < 0.05 with a two-tailed analysis was considered significant).

#### Extracellular Field

##### Slice preparation

Coronal brain slices containing the BLA were collected from 6+ week old male and female mice for *ex vivo* extracellular field recordings. Mice were anesthetized with isoflurane and decapitated using a mouse guillotine. Brains were extracted in ice cold (0 - 4 degree C) sucrose cutting solution containing (in mM) 150 sucrose, 15 glucose, 33 NaCl, 25 NaHCO_3_, 2.5 KCl, 1.25 NaH_2_PO_4_, 1 CaCl_2_, 7 MgCl_2_ (300 - 310 mOsm), oxygenated with 95% O_2_ and 5% CO_2_. Coronal brain slices (350μm) were obtained using a Leica VT1000s vibratome. After cutting, slices were transferred to a petri dish containing sucrose cutting solution where inputs from the hippocampi were removed, and the BLA section placed in an interface incubation chamber filled with continuously oxygenated (95% O_2_, 5% CO_2_) aCSF solution (in mM: 126 NaCl, 10 glucose, 2 MgCl_2_, 2 CaCl_2_, 2.5 KCl, 1.25 NaHCO_3_, 1.5 Na-pyruvate, 1 L-glutamine (300 – 310 mOsm)) set to 34 degrees C. Slices incubated for at least 1 hour before transferring to the recording chamber.

##### Extracellular field recording

BLA sections were placed in an interface recording chamber where a borosilicate glass extracellular field recording electrode was placed in the BLA. LFP data was acquired through LabChart (ADInstruments) at 10 KHz and low pass filtered at 3KHz during acquisition. Gamma oscillations were induced by continuous perfusion of oxygenated modified aCSF with elevated potassium (7.5mM KCl final) and 800nM Kainic acid, at a rate of 1.799mL/min. Baseline and treatment field potentials were recorded for 15 minutes each. For DREADD experiments, only values from the last 5 minutes of each condition were used for analysis. For norepinephrine and WB4101 experiments, values from the last 5 minutes of either baseline or WB4101 were compared against the 3^rd^ to 8^th^ minutes of norepinephrine (roughly capturing the peak NE-induced gamma suppression). All values, regardless of experiment, were normalized to their 15-minute averaged baseline recording. Recording orders are as follows. Norepinephrine experiments: (1) aCSF only, (2) 100μM NE in aCSF. WB4101 experiments: (1) aCSF only, (2) 1μM WB4101 in aCSF, (3) 1μM WB4101 + 100μM NE in aCSF.

### *in vivo* electrophysiology

LFP recordings were performed in awake, freely behaving C57BL/6J and PV-Cre mice, acquired using prefabricated headmounts (Pinnacle Technology, #8201). Frontal cortex EEG and BLA LFP recordings were acquired through a stainless-steel EEG screw and insulated LFP depth electrode implanted over the frontal cortex and in the ipsilateral BLA, respectively. Animals were tethered to the apparatus and EEG and LFP were recorded at 4KHz and amplified 100×. All mice were left to habituate to the recording chamber for at least 30 minutes while tethered before recording.

In PV-Cre animals expressing BLA PV hM3D, baseline and treatment conditions (I.P. injection of saline and CNO (5 mg/kg (1mg/ml), dissolved in saline)) were recorded for 60 min each, and only the last 30 minutes were used for analysis given the IP route of exposure. In cannulated C57 mice, baseline and treatment conditions (intra-BLA infusion of saline, NE, WB4101, and WB4101 + NE) were recorded for 60 minutes each, only the first 30 minutes are shown, and only the first 10 minutes were used for analysis given the infusion route of exposure. All values, regardless of experiment, were normalized to their 60-minute averaged baseline recording.

### Extracellular field data analysis (*in vivo* and *ex vivo*)

LFP and EEG data were band-pass filtered (1-300 Hz, Chebyshev Type II filter), and spectral analysis was performed in MATLAB using publicly-accessible custom-made scripts developed in our lab by Pantelis Antonoudiou (https://github.com/pantelisantonoudiou/MatWAND) utilizing the fast Fourier transform (Frigo & Johnson, 2005). Briefly, recordings were separated into 5 second bins with 50% overlapping segments. The power spectral density for positive frequencies was obtained by applying a Hann window to eliminate spectral leakage. The mains noise (58-60 Hz band) was removed from each bin and replaced using the PCHIP method. Values 3× larger or smaller than the median were considered outliers and replaced with the nearest bin. Processed spectral data were then imported to Python for resampling into one-minute bins and normalization to baseline. Finally, normalized, resampled data were imported to Graphpad Prism for statistical analysis. Statistical comparisons were conducted using either Student’s paired *t* tests or two-way ANOVAs. ANOVA multiple comparisons were corrected for using Tukey’s and Sidak’s multiple comparison tests as appropriate (p < 0.05 was considered significant).

### Cannula fabrication and intra-BLA infusions

Intra-BLA drug infusion cannulas were fabricated in-house using 27G and 30G syringe needles. Twenty-seven gage syringe needles were cut at each end to produce a 10 mm plastic base and 5 mm barrel. Thirty gage syringe needle barrels were cut to 16 mm from the plastic base (including the 1 mm bit of adhesive at the base of the barrel) to create the internal cannula and inserted through the guide cannula to protrude an extra 1 mm. To produce clean syringe barrel openings, barrels were initially cut an extra 1-2 mm longer, and shaved back to the desired length using a Dremel rotary tool (Dremel 4000) with a 120-grit circular sanding attachment.

Intra-BLA infusions were performed using a 25 μL Hamilton syringe affixed to an automated microinfusion pump (Harvard Apparatus, The Pump 11 Elite Nanomite), and connected to the internal cannula needle via plastic tubing (Tygon flexible plastic tubing; ID = 0.020 IN, OD = 0.060 IN). Intra-BLA infusions of 300 nL norepinephrine (10 mM), WB4101 (10 μM), or saline were administered at a rate of 0.2 μL/min. After infusion, the needle was left to sit for an extra minute past the infusion to allow for sufficient diffusion and minimize backflow upon removal from the brain.

