## Supplementary Figures for "Neuromodulation-induced burst firing in parvalbumin interneurons of the basolateral amygdala mediates transition between fear-associated network and behavioral states"

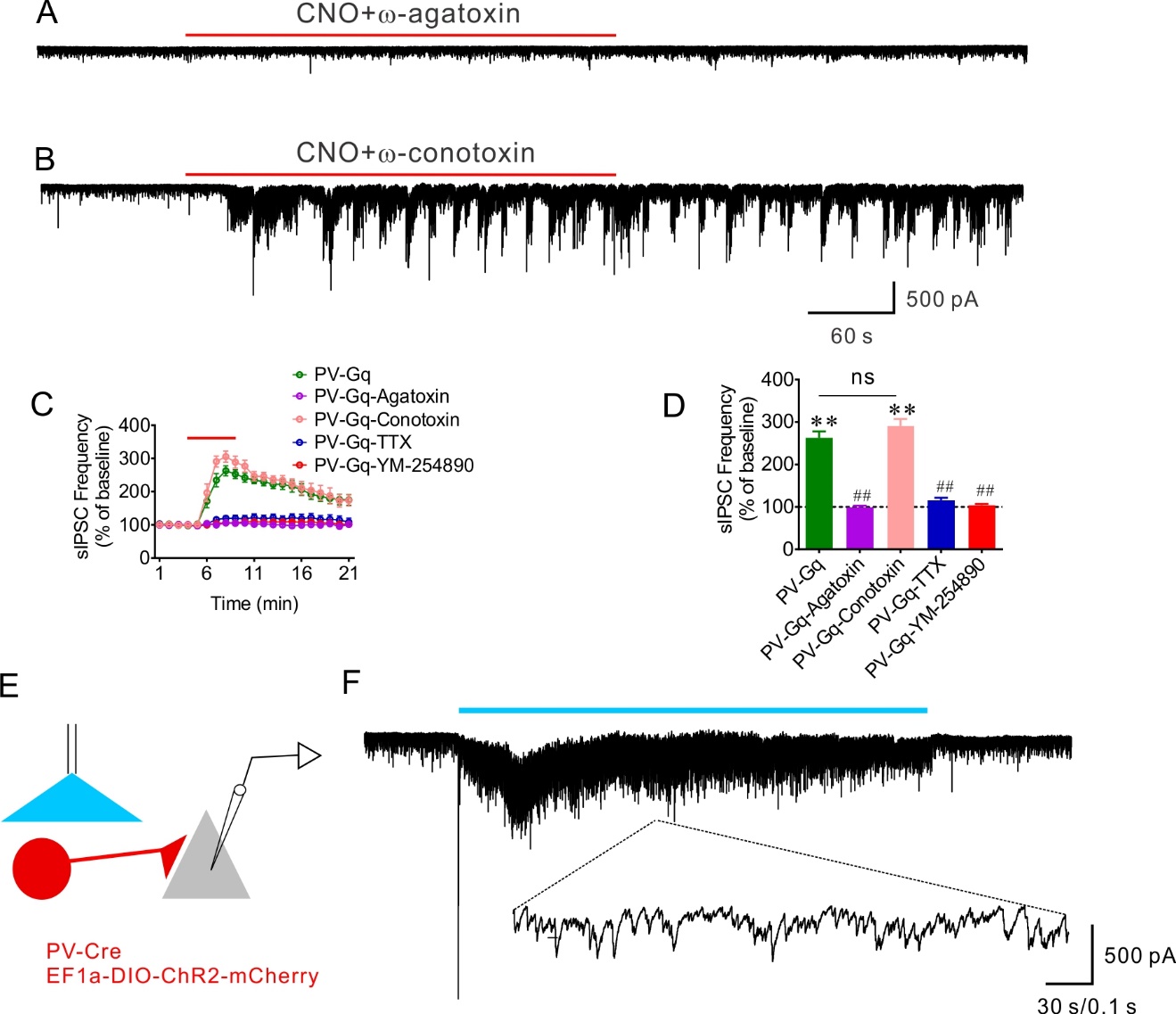

Supplementary Figure 1 (Supporting Figure 1). (A, B) Representative recordings of PV interneuron-mediated repetitive IPSC bursts in a BLA principal neuron, which were blocked by pre-treatment of the slice with the P/Q-type calcium channel blocker ω-agatoxin, but were unaffected by pre-incubation of the slice in the N-type calcium channel blocker ω-conotoxin. (C, D) Time course and mean change in sIPSC frequency in response to Gq activation in PV interneurons, which was blocked by ω-agatoxin and insensitive to ω-conotoxin. Pre-treatment of the slice with TTX and YM-254890 abolished the effect of Gq activation in PV neurons on sIPSC frequency. (PV-Gq, n=12 cells from 5 mice; PV-Gq-Agatoxin, n=7 cells from 3 mice; PV-Gq-Conotoxin, n=7 cells from 3 mice; PV-Gq-TTX, n=8 cells from 4 mice; PV-Gq-YM-254890, n=9 cells from 4 mice) (Paired *t* test, PV-Gq vs. baseline: p < 0.0001, PV-Gq-Conotoxin vs. baseline: p < 0.0001, ** p < 0.01; One-Way ANOVA, F (4,38) = 63.35, p < 0.0001; Dunnett’s multiple comparisons test, PV-Gq vs. PV-Gq-Agatoxin: p < 0.0001, PV-Gq vs. PV-Gq-Conotoxin: p = 0.27, PV-Gq vs. PV-Gq-TTX: p < 0.0001, PV-Gq vs. PV-Gq-YM-254890: p < 0.0001, ## p < 0.01, ns not significant). (E) Schematic diagram of recording from BLA principal neurons after transduction and light activation of ChR2 in BLA PV interneurons. (F) Representative recording of the response of a BLA principal neuron to activation of PV interneurons via continuous photostimulation, which generated a tonic increase in IPSCs. Bottom: Expanded trace showing the tonic generation of IPSCs.

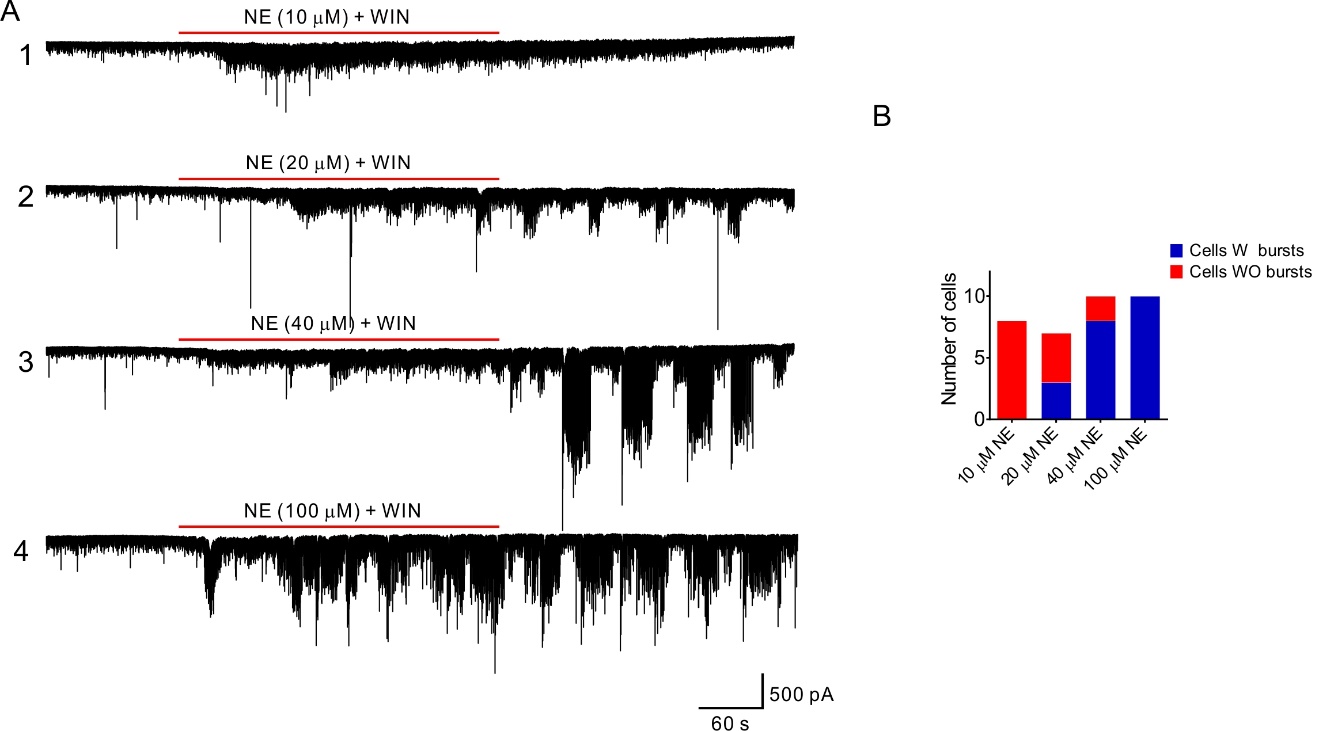
Supplementary Figure 2 (supporting Figure 2). Norepinephrine concentration dependence of sIPSC bursting in BLA principal cells. (A) Representative traces showing the IPSC responses in BLA principal neurons to 1) 10 μM, 2) 20 μM , 3) 40 μM, and 4) 100 μM NE application. The CB1 receptor agonist WIN 55,212-2 (1 μM) was applied to suppress CCK neuron-mediated IPSCs. At concentrations ≥ 20 μM, NE induced repetitive bursts of IPSCs in BLA principal cells. (B) Quantification of the proportion of recorded cells that generated bursts of IPSCs at the different NE concentrations.

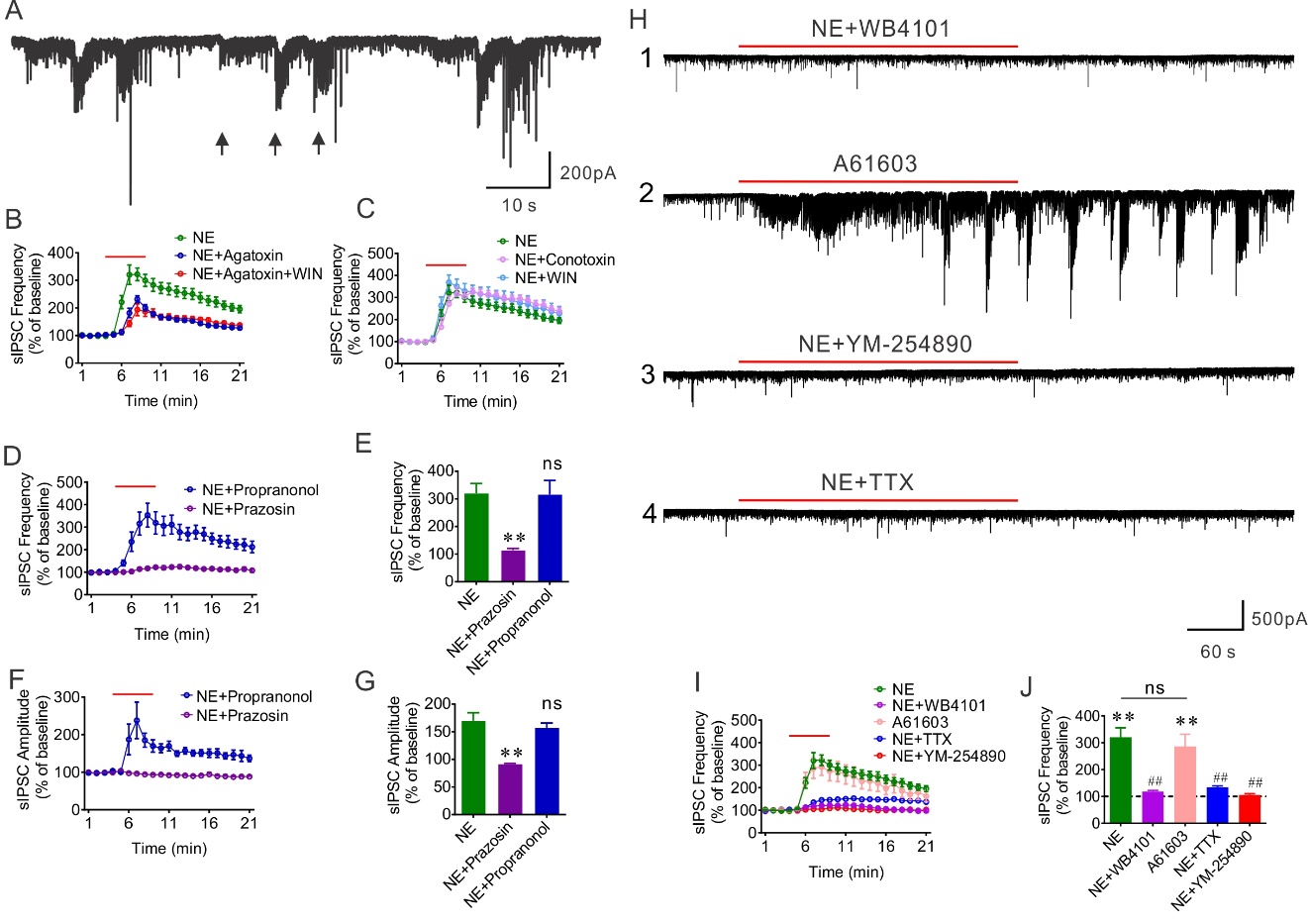
Supplementary Figure 3 (supporting Figure 3). Alpha 1-adrenergic increase in sIPSC frequency and amplitude mediated by presynaptic PV neuron activation. (A) A representative recording of multiple types of NE-induced repetitive IPSC bursts (arrows) in a BLA principal neuron. (B) Time course of the NE-induced change in sIPSC frequency with and without blockade of P/Q-type calcium channels (ω-agatoxin) and activation of CB1 receptors (WIN). Blocking P/Q-type calcium channels suppressed the NE-induced increase in IPSC frequency, while the residual frequency response was not further blocked by CB1 activation. (C) Time course of the effect of blocking N-type calcium channels and activating CB1 receptors on sIPSC frequency. Neither treatment had any effect on the NE-induced increase in sIPSC frequency, indicating a lack of contribution of CCK interneurons. (D, E) Time course and mean change in sIPSC frequency showing that the NE-stimulated IPSCs are blocked by the broad-spectrum α1 adrenoreceptor antagonist prazosin, but are unaffected by the β adrenoceptor antagonist propranolol. (NE: 16 cells from 5 mice; NE + Prazosin: 6 cells from 4 animals; NE + Propranolol: 8 cells from 3 mice) (One-Way ANOVA, F (2, 27) = 6.07, p = 0.0066; Dunnett’s multiple comparisons test, NE vs. NE + Prazosin: p = 0.0046, NE vs. NE + Propranolol: p = 0.99, **, p < 0.01, ns, not significant). (F, G) Time course and mean change in sIPSC amplitude in response to NE in the presence of prazosin and propranolol. Blocking α1 receptors with prazosin abolished the NE effect, but blocking β receptors with propranolol had no effect on the NE facilitation of sIPSC amplitude. (NE: 16 cells from 5 mice; NE + Prazosin: 6 cells from 4 animals; NE + Propranolol: 8 cells from 3 mice) (One-Way ANOVA, F (2, 27) = 6.25, p = 0.0059, Dunnett’s multiple comparisons test, NE vs. NE + Prazosin: p = 0.0031, NE vs. NE + Propranolol: p = 0.75, **, p < 0.01, ns, not significant). (H) Representative recordings showing that the NE-induced facilitation of sIPSCs depends on activation of presynaptic interneurons through Gq-coupled α1A adrenoreceptors. (1) the selective α1A adrenoreceptor antagonist WB4101 blocked the NE-stimulated sIPSCs. (2) the α1A adrenoreceptor agonist A61603 induced repetitive bursts of IPSCs following the initial increase of IPSCs, similar to the effect of NE. (3) Blocking Gq activity with YM-254890 eliminated the NE-induced increase in sIPSCs. (4) Blocking spike activation with TTX prevented the NE facilitation of sIPSCs. (I, J) Time course and mean change in sIPSC frequency. The NE-induced increase in sIPSC frequency was blocked by the α1A adrenoceptor antagonist WB4101 and was mimicked by the α1A adrenoceptor agonist A61603. Blocking either spiking activity with TTX or Gq activation with YM-254890 eliminated NE induced increase in sIPSC frequency. (NE: 16 cells from 5 mice; NE + WB4101: 7 cells from 4 mice; A61603: 10 cells from 4 mice; NE + TTX: 10 cells from 5 mice; NE + YM-254890: 8 cells from 3 mice) (paired *t* test, NE vs. baseline, p < 0.0001, A61603 vs. baseline: p = 0.0027, ** p < 0.01; one-way ANOVA, F (4, 46) = 10.73, p < 0.0001, Dunnett’s multiple comparisons test, NE vs. NE + WB4101: p = 0.0003, NE vs. A61603: p = 0.83, NE vs. NE + TTX: p = 0.0002, NE vs. NE + YM-254890: p < 0.0001, ##, p < 0.01, ns, not significant).

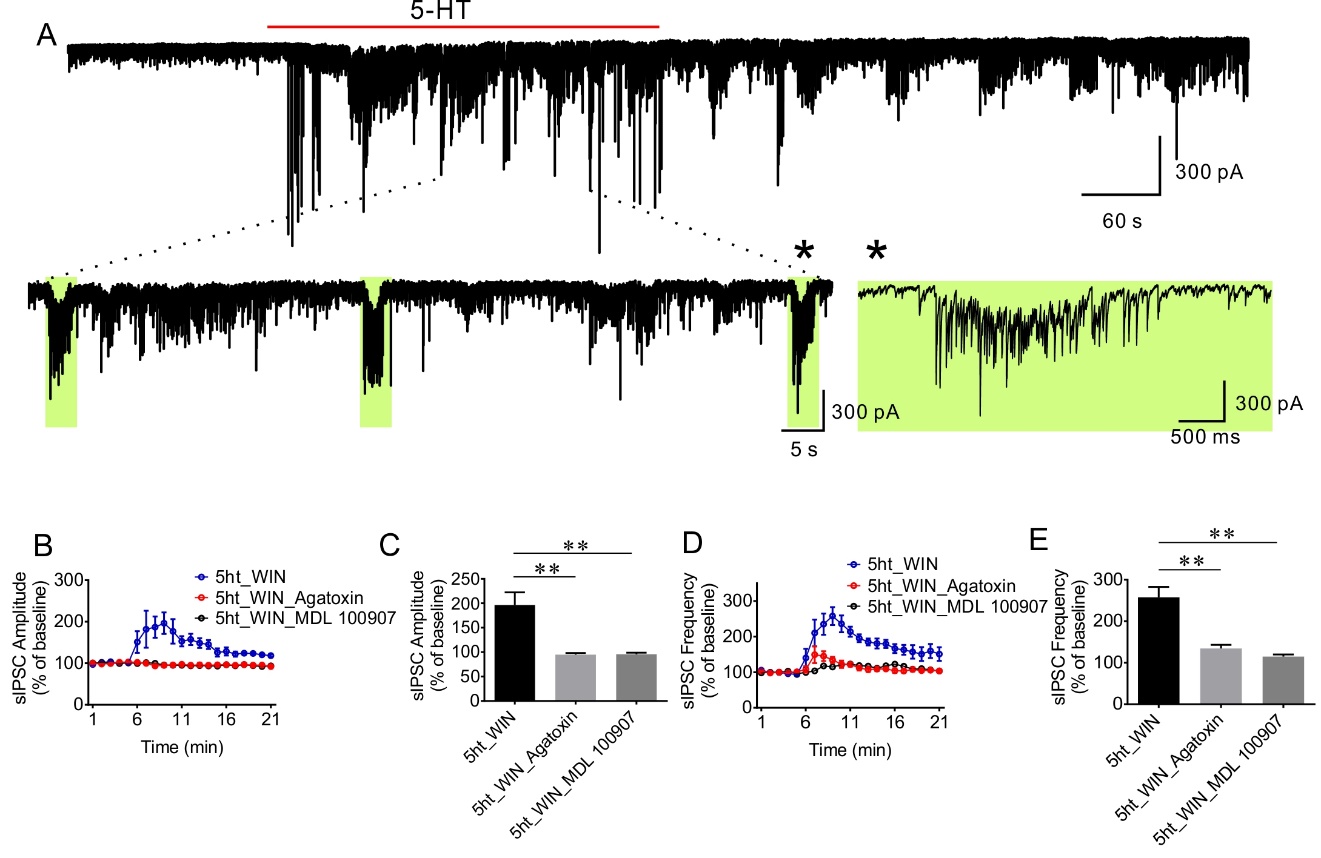

Supplementary Figure 4 (supporting Figure 3). Serotonin 5-HT2A receptor activation of IPSC bursts. (A) A representative recording of serotonin (5-HT, 100 µM)-induced repetitive IPSC bursts in a BLA principal neuron in the presence of glutamate receptor antagonists, DNQX and AP5, and the CB1 receptor agonist WIN 55,212-2. Green boxes show three repetitions of the 5-HT-induced bursts. The last repetition was expanded to show the accelerating frequency of intra-burst IPSCs and the negative shift in the baseline due to the progressive summation of high-frequency IPSCs. (B, C) Time course and mean change in sIPSC amplitude in response to 5-HT. The 5-HT-induced IPSC bursts were blocked by the P/Q type calcium channel blocker, ω-agatoxin, and the Gq-coupled 5HT2A receptor antagonist, MDL 100907 (1 µM) (5-HT + WIN, n=5 cells from 3 mice; 5-HT + WIN + agatoxin, n=5 cells from 3 animals; 5-HT + WIN + MDL 100907, n=5 cells from 2 mice; one-way ANOVA, F (2, 12) = 14.77, p = 0.0006, Dunnett’s multiple comparisons test, 5-HT + WIN vs. 5-HT + WIN + agatoxin: p = 0.0009, 5-HT + WIN vs. 5-HT + WIN + MDL 100907: p = 0.001, **, p < 0.01). (D, E) Time course and mean change in sIPSC frequency in 5-HT. Similar to NE, the 5-HT-induced IPSC bursts were blocked by the P/Q blocker ω-agatoxin and the 5HT2A receptor MDL 100907. (5-HT+WIN, n=5 cells from 3 mice; 5-HT + WIN + agatoxin, n=5 cells from 3 animals; 5-HT + WIN + MDL 100907, n=5 cells from 2 mice; One-Way ANOVA, F (2, 12) = 24.86, p < 0.0001, Dunnett’s multiple comparisons test, 5-HT + WIN vs. 5-HT + WIN + agatoxin: p = 0.0002, 5-HT + WIN vs. 5-HT + WIN + MDL 100907: p < 0.0001, ** p<0.01).

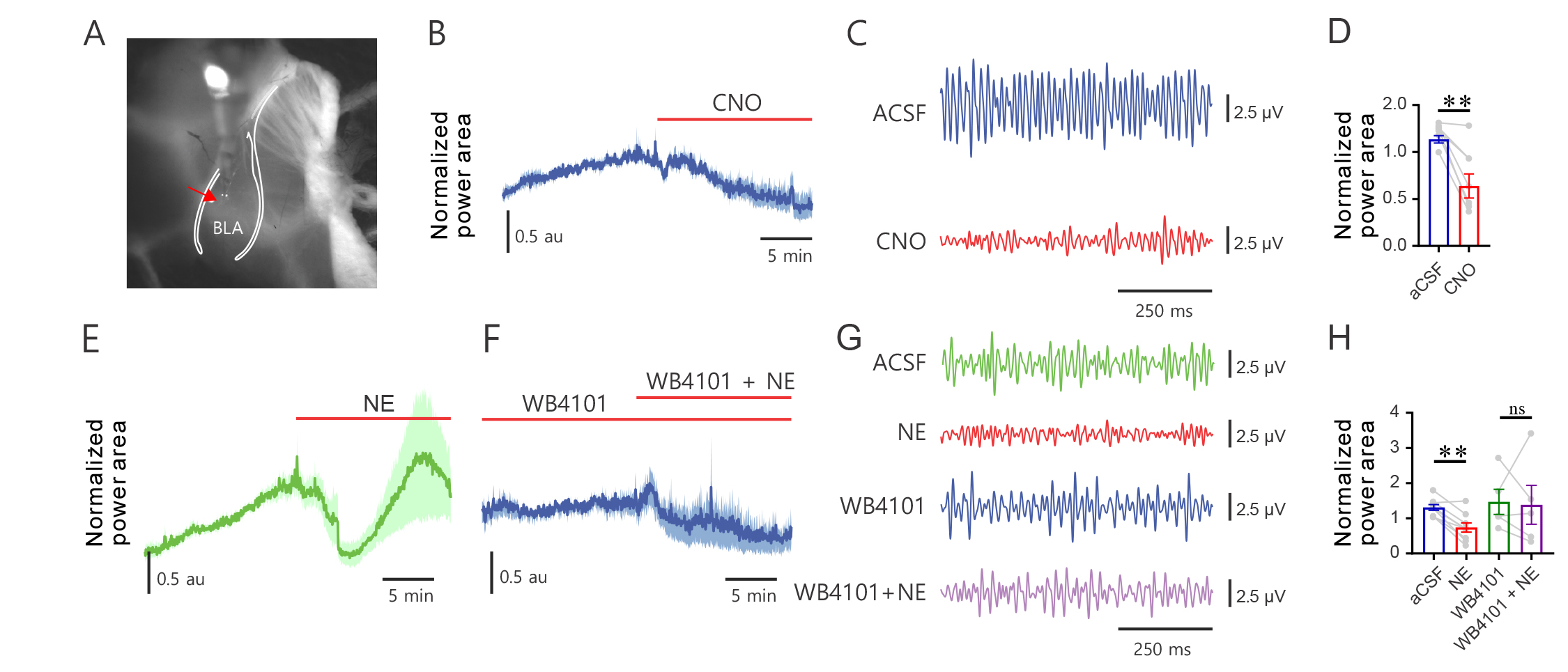
Supplementary Figure 5 (supporting Figure 6). PV Gq signaling reconfigures BLA network activity *ex vivo*. (A) Phase contrast image of the resected BLA slice preparation. Arrow denotes electrode placement. (B) Normalized gamma power (30-80 Hz) area over time before and during CNO activation of virally transduced hM3D-expressing PV neurons in BLA slices from PV-Cre mice. (C) Bandpass filtered (30-80 Hz) raw traces during ACSF (top) or CNO (bottom) application. (D) Mean (+/- SEM) normalized gamma power area across treatments. Paired *t* test, ACSF vs CNO (n=7 slices from 4 animals, p = 0.0036). Values from 5 min before and the last 5 min of CNO treatment were used for analysis. **, p < 0.01 (E-F) Normalized gamma power area over time across conditions, (E) ACSF, NE, (F) WB4101, WB4101 + NE. (G) Bandpass filtered (30-80 Hz) raw traces across treatments (top-bottom: ACSF, NE, WB4101, WB4101+NE). (H) Mean (+/- SEM) normalized power area across treatments. Paired *t* test, ACSF vs NE (n=9 slices from 4 animals, p = 0.0003); WB4101 vs WB4101+NE (n=5 slices from 3 animals, p = 0.8706). Values from minutes 3 to 8 were used for analysis (capturing the peak NE-induced gamma suppression). *, p < 0.01, ns = not significant, au = arbitrary units.

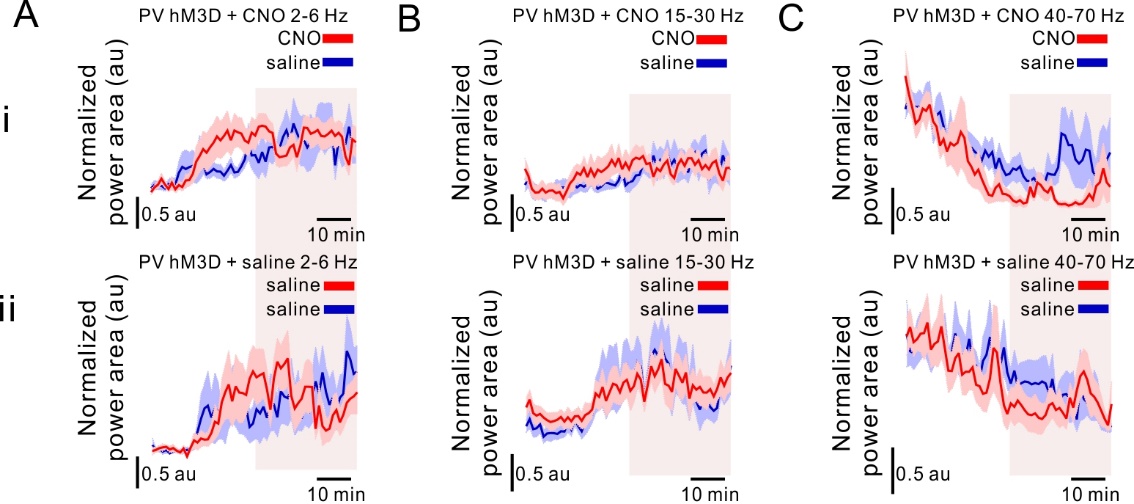

Supplementary Figure 6 (supporting Figure 6). Effect of hM3D-Gq activation in PV neurons on normalized power area timeseries in nonsignificant frequency bands. (A-C) Normalized power area over time across frequency bands (A) 2-6 Hz, (B) 15-30 Hz, (C) 40-70 Hz. (i-ii) Normalized power area over time across treatments (i) CNO (5 mg/kg IP) vs Saline, (ii) Saline1 vs Saline2. Highlighted sections denote values included for analysis in Figure 7. Values averaged in 1-min bins. Colored lines = average power over time, colored shaded areas = SEM. au = arbitrary units.

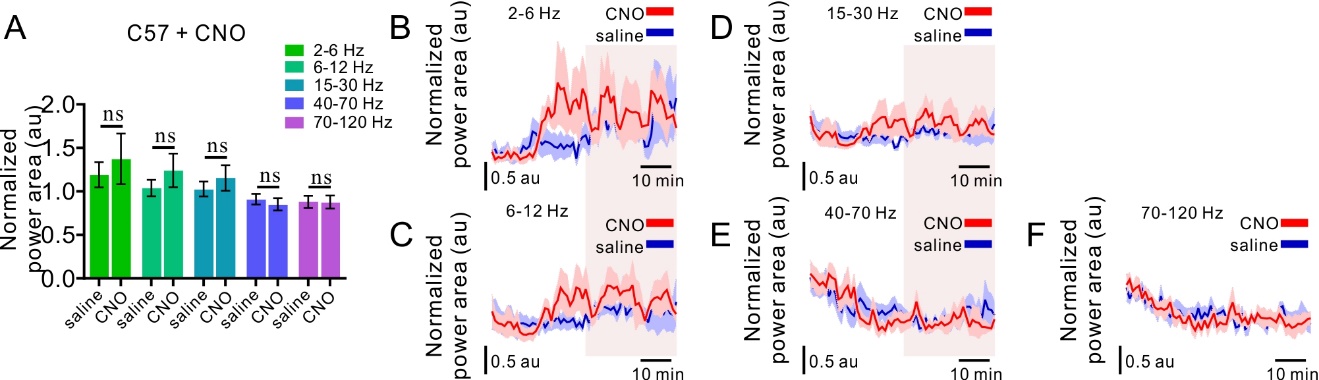

Supplementary Figure 7 (supporting Figure 6). CNO does not significantly modulate BLA oscillatory activity in control C57BL/6J mice. (A) Mean (+/- SEM) normalized power area across treatments (CNO vs Saline) and frequency bands. Values from last 30 minutes of recording were used for analysis, illustrated by the highlighted sections in B-F. Two-way ANOVA [treatment x frequency], F (4, 16) = 1.702, p = 0.1987. Two-way ANOVA [treatment], F (1, 4) = 3.405, p = 0.1388. (B-D) Normalized power area across time for (B) 2-6 Hz, (C) 6-12 Hz, (D) 15-30 Hz, (E) 40-70 Hz, (F) 70-120 Hz. Values averaged into 1-minute bins. Line = average power over time, shaded area = SEM. n = 5. ns = not significant, au = arbitrary units.

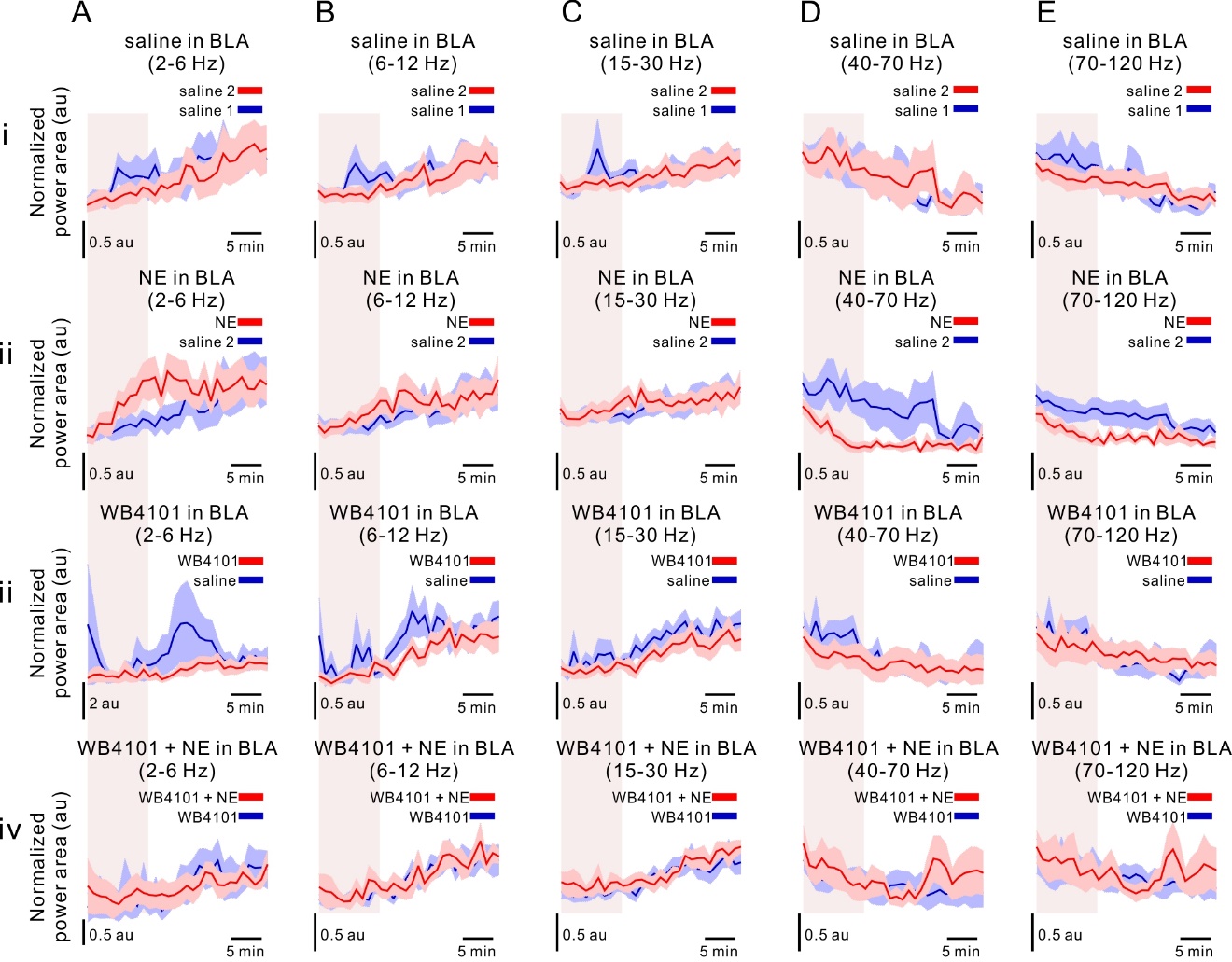

Supplementary Figure 8 (supporting Figure 7). Effect of intra-BLA α1A adrenoreceptor activation on BLA LFP oscillations. Saline/WB4101/NE/WB4101+NE timeseries data. (A-C) Normalized power area over time across frequency bands (A) 2-6 Hz, (B) 6-12 Hz, (C) 15-30 Hz, (D) 40-70 Hz, (E) 70-120 Hz. (i-iv) Normalized power area over time across treatments (i) Saline1 vs Saline2, (ii) Saline2 vs NE, (iii) Saline vs WB4101, (iv) WB4101 vs WB4101 + NE. Highlighted sections denote values included for analysis in figure 8. Values averaged into 1-minute bins. Line = average power over time, shaded area = SEM. au = arbitrary units.

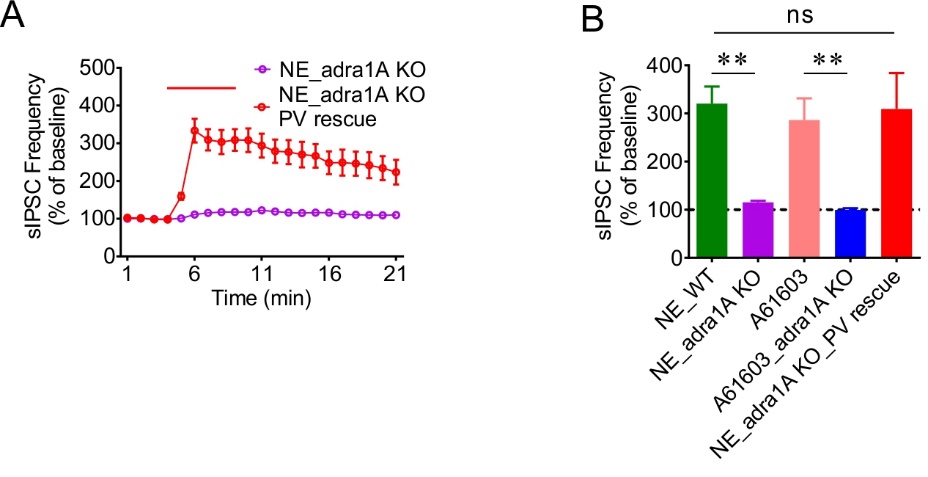
Supplementary Figure 9 (supporting Figure 8). Effect of adra1A knockout and rescue on NE-induced increase in sIPSC frequency. (A) Time course of NE effect of sIPSC frequency in adra1A KO mice with or without re-expression of α1A adrenoreceptors in BLA PV interneurons. (B) Mean change in sIPSC frequency showing that rescue of α1A noradrenergic signaling in BLA PV interneurons restored the NE facilitation of sIPSC frequency that was largely lost in BLA principal neurons in slices from adra1A KO mice (NE_WT: 16 cells from 5 mice; NE_adra1A KO: 9 cells from 4 mice; A61603_WT: 10 cells from 4 mice; A 61603_adra1A KO: 7 cells from 3 mice; NE_adra1A KO_PV rescue: 7 cells from 3 mice) (unpaired *t* test, A61603_WT vs. A61603_adra1A KO: p=0.0037; One-Way ANOVA, F (2, 29) = 11.43, p = 0.0002, Dunnett’s multiple comparisons test, NE_WT vs. NE_adra1A KO: p = 0.0002, NE_WT vs. NE_adra1A KO_PV rescue: p = 0.96, ** p < 0.01, ns, not significant).

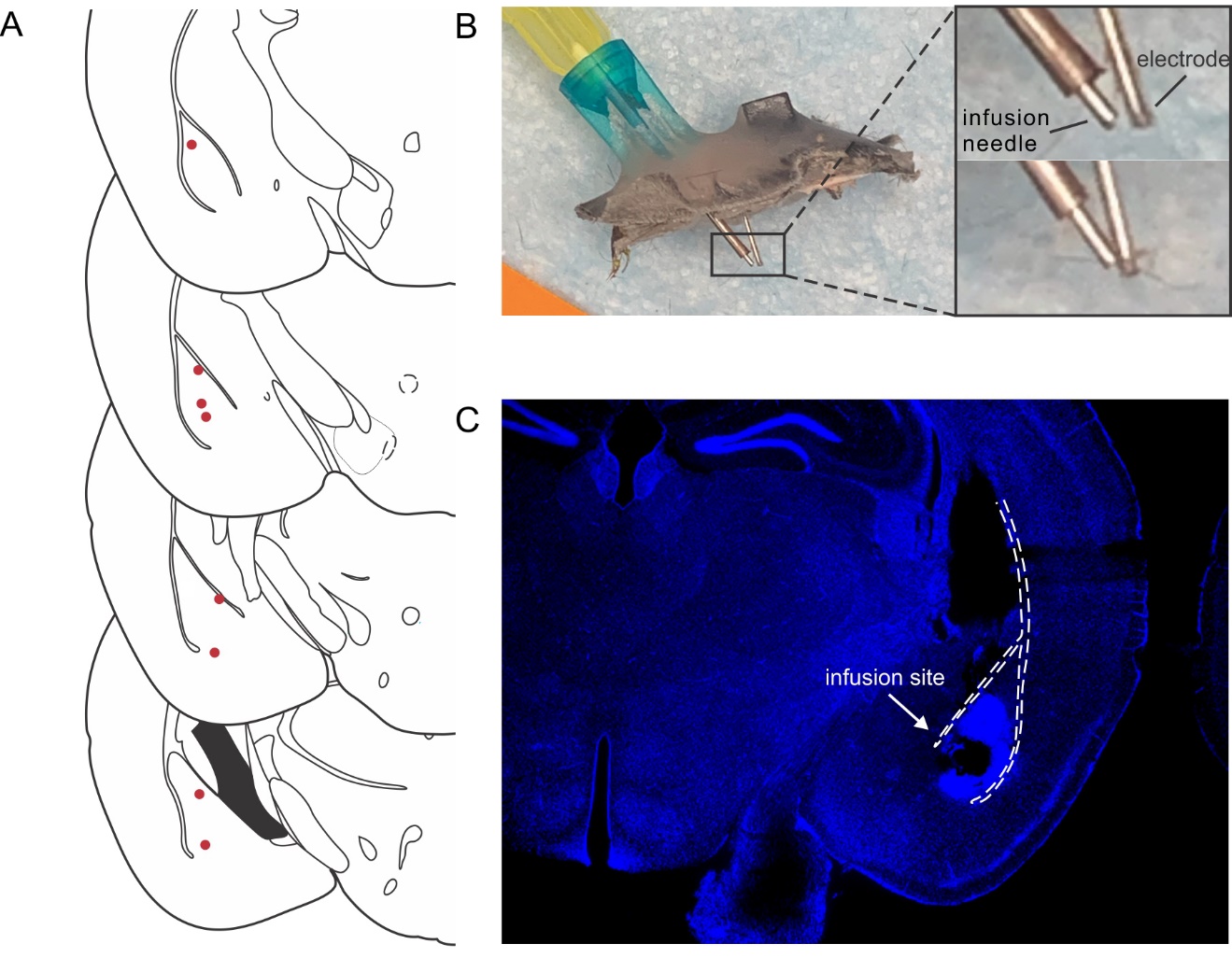

Supplementary Figure 10. Intra-BLA infusion cannula placement. (A) BLA cannula placement across animals. (B) Extracted representative implant showing electrode placement relative to microinfusion cannula. Right: higher magnification images of two implants illustrating the variable distance between the microinfusion cannula and LFP recording electrode. (C) Coronal BLA section demonstrating BLA cannula placement accuracy. The lesioned area indicates infusion site within the BLA.
